# Leukocyte dynamics after intracerebral hemorrhage in a living patient reveal rapid adaptations to tissue milieu

**DOI:** 10.1101/2020.11.10.375675

**Authors:** Brittany A. Goods, Michael H. Askenase, Erica Markarian, Hannah E. Beatty, Riley Drake, Ira Fleming, Jonathan H. DeLong, Naomi H. Philip, Charles C. Matouk, Issam A. Awad, Mario Zuccarello, Daniel F. Hanley, J. Christopher Love, Alex K. Shalek, Lauren H. Sansing, on behalf of the ICHseq Investigators

**Author notes:** equal (Brittany A. Goods, Michael H. Askenase). equal (J. Christopher Love, Alex K. Shalek, Lauren H. Sansing). Corresponding Authors: Lauren Sansing, Alex K. Shalek, J. Christopher Love.

## Abstract

Intracerebral hemorrhage (ICH) is a devastating form of stroke with a high mortality rate and few treatment options. Discovery of therapeutic interventions has been slow given the challenges associated with studying acute injury, particularly over time, in the human brain. Inflammation induced by exposure of brain tissue to blood appears to be a major part of brain tissue injury. Here we longitudinally profiled blood and cerebral hematoma effluent from a patient enrolled in the Minimally Invasive Surgery with Thrombolysis in Intracerebral Haemorrhage Evacuation (MISTIEIII) trial, offering a rare window into the local and systemic immune responses to acute brain injury. Using single-cell RNA-sequencing, we characterized the local cellular response during ICH in the brain of a living patient at single-cell resolution for the first time. Our analysis revealed rapid shifts in the activation states of myeloid and T cells in the brain over time, suggesting that leukocyte responses are dynamically reshaped by the hematoma microenvironment. Interestingly, the patient had an asymptomatic re-bleed (second local exposure to blood) that our transcriptional data indicated occurred more than 30 hours prior to detection by CT scan. This case highlights the rapid immune dynamics in the brain after ICH and suggests that sensitive methods like scRNA-seq can inform our understanding of complex intracerebral events.

## INTRODUCTION

Intracerebral hemorrhage (ICH), brain bleeding often caused by hypertension, has a worldwide incidence of over 3 million cases per year, limited treatment options, and high mortality rate (40-60% worldwide).^1,2^ To date, surgical approaches to remove the hemorrhage have not improved functional outcomes.^3,4^ The Phase III clinical trial of Minimally Invasive Surgery with Thrombolysis in Intracerebral Hemorrhage Evacuation (MISTIE III) was designed to test whether minimally invasive catheter evacuation followed by thrombolysis would improve functional outcome in patients with ICH. The trial’s approach used image-guided minimally invasive surgery to aspirate the liquid component of the hemorrhage followed by placement of a sutured indwelling catheter to liquify and drain remaining hemorrhage over several days with the aid of recombinant tissue plasminogen activator (tPA). The trial found that surgery was safe and reduced mortality, but did not improve overall functional outcomes 365 days after ICH.^5^ As part of a trial sub-study (ICHseq), daily samples of both hematoma effluent from the catheter and peripheral blood were collected. These samples provided a unique opportunity to study the immune response to ICH over time in living patients.

Here, we describe longitudinal cellular characterization of hematoma effluent and blood from a patient who was enrolled in MISTIE III using single-cell RNA-sequencing (scRNA-seq).^6^ We define distinct leukocyte phenotypes that may dynamically reflect local inflammatory responses in the brain as well as the emergence of cells resembling peripheral blood leukocytes that we propose identify the onset of repeat tissue exposure to blood (in this case, an asymptomatic re-bleeding event) later detected by CT scan. Additionally, we show that leukocyte transcriptional programs in the patient’s blood return to a baseline comparable to age-matched control blood 2.5 years after ICH. This case report is the first longitudinal single-cell genomic characterization of cellular infiltrate in acute brain injury in a living patient and, importantly, underscores the utility of single-cell analyses for understanding dynamic immune responses in the brain and other tissues.

## RESULTS AND DISCUSSION

A 74-year old female with hypertension presented to the emergency room with difficulty speaking and right-sided weakness (**Supplemental Table 1)**. Her initial National Institutes of Health Stroke Scale Score (NIHSS) was a 19, and symptoms included aphasia and right-sided hemiplegia, sensory loss, and visual field deficit. A CT scan of the brain showed a large left ICH. Two days after ICH onset, she was enrolled into the MISTIE III trial and randomized to the surgical arm. At 51 hours post-hemorrhage, 45 mL of liquid hematoma were aspirated and the drainage catheter was placed. The following day, instillation of tPA (1 mg every 8 hours) was initiated per trial protocol with subsequent effective drainage of the remaining hematoma (**Figure 1A**). Beginning on day three post-hemorrhage, an increasing volume of drainage was noted from the catheter and tPA was discontinued. The patient consistently improved neurologically. The catheter continued to drain the hematoma for two more days before being removed. The patient was subsequently discharged to an acute rehabilitation facility. Thirty days after ICH onset, she had improved to mild aphasia (difficulty speaking), right visual field deficit, and mild right hemiparesis (weakness) and continued to recover functionally. At 2.5 years after onset, she was active with minimal aphasia and right-sided vision loss (NIHSS 3).

**Figure 1.**
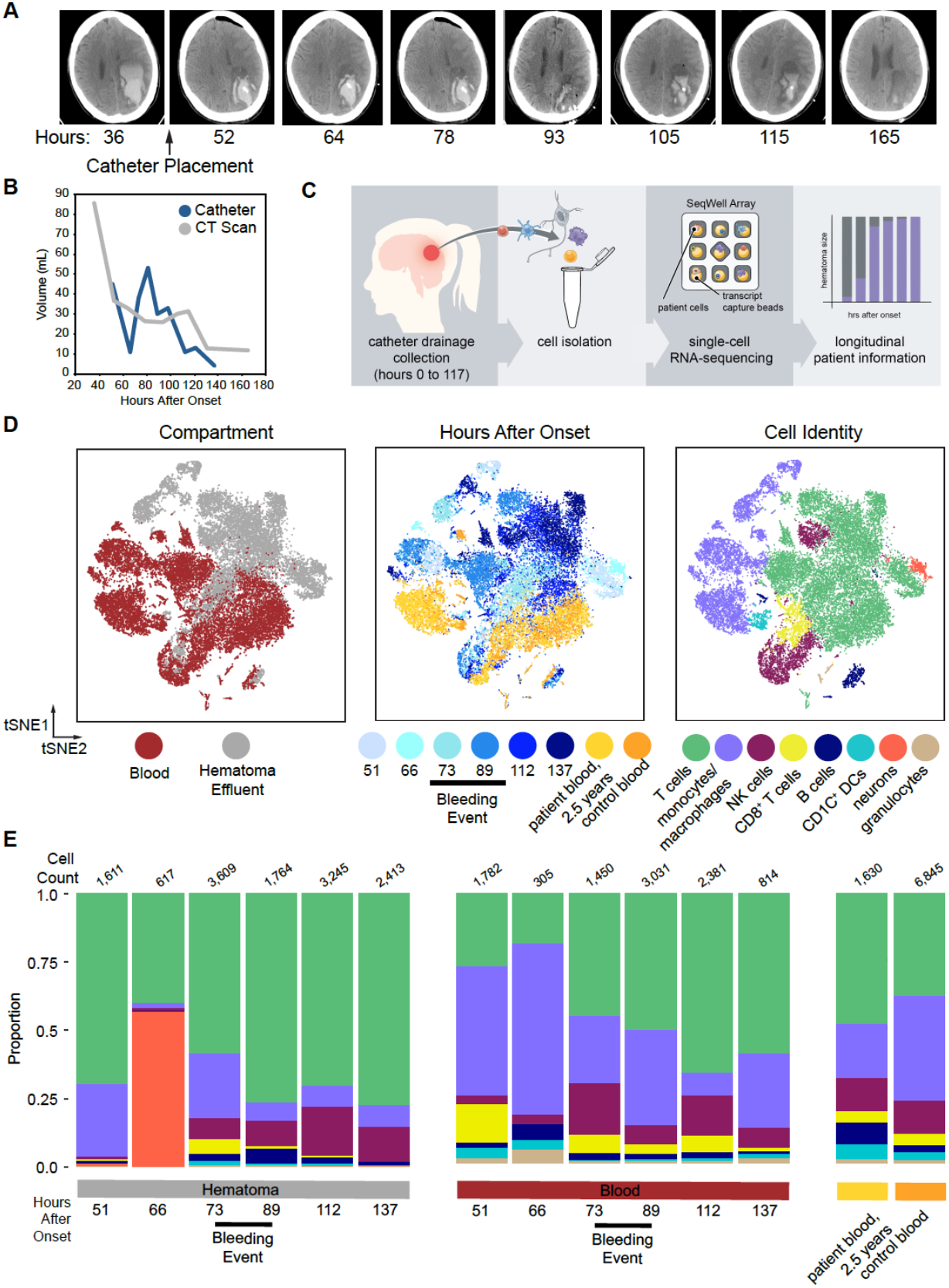
scRNA-seq on cells isolated from hematoma effluent and peripheral blood in a living patient over the course of ICH. **A**. Selected CT scans as a function of time after onset (hrs). The time of catheter placement is indicated. **B**. Plot of hematoma volume and drainage as a function of time as measured by both CT scan and the volume of hematoma effluent from the catheter in the eight hours prior to collection. **C**. Schematic overview of sample collection and processing for generating scRNA-seq data from patient hematoma effluent and blood. Single cells from hematoma effluent and blood were isolated, and scRNA-seq profiles were generated using the Seq-Well platform. **D**. t-distributed stochastic neighbor embedding (tSNE) plot along components 1 and 2 for all high-quality single cells (n=24,877 single cells across seven patient time points and n=6,845 single cells across four control donors). The tSNE plot is colored by compartment of origin (left), time after onset (middle), or cell identity (right). Cell identity was determined by SNN clustering, marker selection, and module scoring (**Supplemental Figure 2, Supplemental Methods**). The estimated onset of the asymptomatic re-bleeding event is indicated with a black bar and is based on changes detected in CT scan, catheter drainage data, and sequencing data. **E**. Stacked frequency plots for each indicated condition and colored by identified cell type. Estimated earliest time point of re-bleeding event is indicated. Total cell counts per cluster are reported directly in **Supplemental Table 3**.

Overall, during the patient’s hospital stay, her hematoma was effectively drained (**Figure 1A**) and she improved clinically. Midway through her treatment increasing hematoma volume by CT scan and an increase in hematoma drainage suggested that she experienced an asymptomatic re-bleeding event and treatment with tPA was halted (**Figure 1A, Supplemental Table 1**). Volumetric quantification by serial CT scans suggests that this event occurred between 93 and 105 hours after the onset of the hemorrhage. Increases to the volume and cellularity of hematoma drainage, however, were observed prior to hematoma expansion as measured by imaging, beginning to rise by 73 hours after onset **(Figure 1B, Supplemental Table 1)**. These data suggest that the patient’s re-bleeding event occurred prior to the increased volume seen on CT scan.

We hypothesized that detailed analysis of the cellular composition of the hematoma effluent in this patient would provide insight into the molecular features of the hematoma microenvironment, as well as her re-bleeding event, and determine whether immune responses are brain tissue-specific. Using Seq-Well, we performed scRNA-seq on individual hematoma cells and peripheral blood collected longitudinally during her hospitalization as well as peripheral blood collected after 2.5 years, and control blood from age-matched healthy donors (**Figure 1C**).^6^ We generated single-cell transcriptional profiles for 31,722 cells spanning seven time points after onset, and these are displayed along with control samples using a t-distributed stochastic neighbor embedding (tSNE) (**Figure 1D, Supplemental Figure 1 and 2**) along the first two components. Replicates (performed for most time points) clustered together across each time point and have comparable quality control metrics (**Supplemental Figure 1**). Overall, we found that blood and hematoma-derived cells were transcriptionally distinct at almost all time points, and separated in tSNE space by compartment, suggesting that the local milieu within the hematoma is a major driver of leukocyte transcriptional state and function after ICH.

After removing red blood cells (**Supplemental Methods**), we identified seven major cell types, including neurons, B cells, T cells, monocytes, dendritic cells, NK cells and neutrophils (**Figure 1D, Supplemental Figure 3A, B, and C**). We found that monocytes/macrophages and T cells were the most abundant in both peripheral blood and hematoma samples. The other major immune cell types were consistently identified in every sample, were consistent across replicate samples and were comparable in quality (**Supplemental Figure 3B and D**). We confirmed these findings using separately collected longitudinal samples collected from this patient and analyzed by mass cytometry (**Supplemental Figure 4, Supplemental Table 2**). Our mass cytometry results further corroborate our scRNA-seq frequencies, showing that T cells and monocytes were the predominant cell types profiled in both blood and hematoma. Finally, in our scRNA-seq data, we found a large cluster of neurons that appeared exclusively in hematoma effluent at high frequency prior to increased hematoma drainage at 66 hours (**Figure 1D and E, Supplemental Table 2**). These neurons were the sole cluster positive for KIF5A, a neuron-specific kinesin subunit (**Supplemental Figure 3C**).^7,8^

To assess whether the leukocyte populations identified in the peripheral blood after ICH differed from baseline, we also generated single-cell transcriptional profiles from the patient’s blood collected 2.5 years post-ICH and from the blood of four age-matched healthy donors (**Figure 1D and E**). The follow-up blood and healthy control blood generally overlap, and may suggest an overall return to baseline in circulating leukocytes in the blood of this patient (**Supplemental Figure 2**). Bulk RNA-seq analysis on sorted populations of immune cells, including CD4^+^ T cells and monocytes (**Supplemental Figure 5 A and B**), from this patient’s blood and hematoma at several time points (**Supplemental Table 1**), combined with hallmark gene set enrichment analysis (**Supplemental Figure 5C**), corroborates the substantial transcriptomic variation over time despite little variability in broad immune cell frequencies assessed by flow cytometry (data not shown).

In the MISTIE III trial, 2% of patients in the surgical arm had a symptomatic re-bleeding event, however 32% had asymptomatic re-bleeding.^5,9^ Hematoma effluent volume drainage is an imperfect measurement of rebleeding, as some hematoma cavities have variable connections with the ventricular or subarachnoid space and, at times, volume may increase due to CSF drainage. This event is difficult to identify or predict at the bedside. The implications of rebleeding on the local immune response and long-term outcomes remain unknown but have an important impact of treatment (i.e., cessation of tPA). We hypothesized that a deeper analysis of the predominant cell types in our scRNA-seq data, including T cells and myeloid cells, could better delineate the compartment-specific response to an asymptomatic re-bleed. To accomplish this, we performed sub-clustering analyses on each population separately (**Supplemental Methods**).

Among myeloid cells (macrophage, monocyte/macrophage, and dendritic cells),^10^ we found that separation by compartment (i.e., blood versus hematoma) was preserved (**Figure 2A**) and identified 14 sub-clusters, as well as defining genes for each (**Figure 2A, Supplemental Figure 6A, Supplemental Table 4**). Three of these (myeloid sub-clusters 2, 4, and 5) were almost exclusively found in hematoma effluent (**Figure 2B, Supplemental Figure 6B**). We found that the frequency of myeloid sub-clusters shifted over time, with several appearing in either blood or hematoma around the time of re-bleeding (**Figure 2B**).

**Figure 2.**
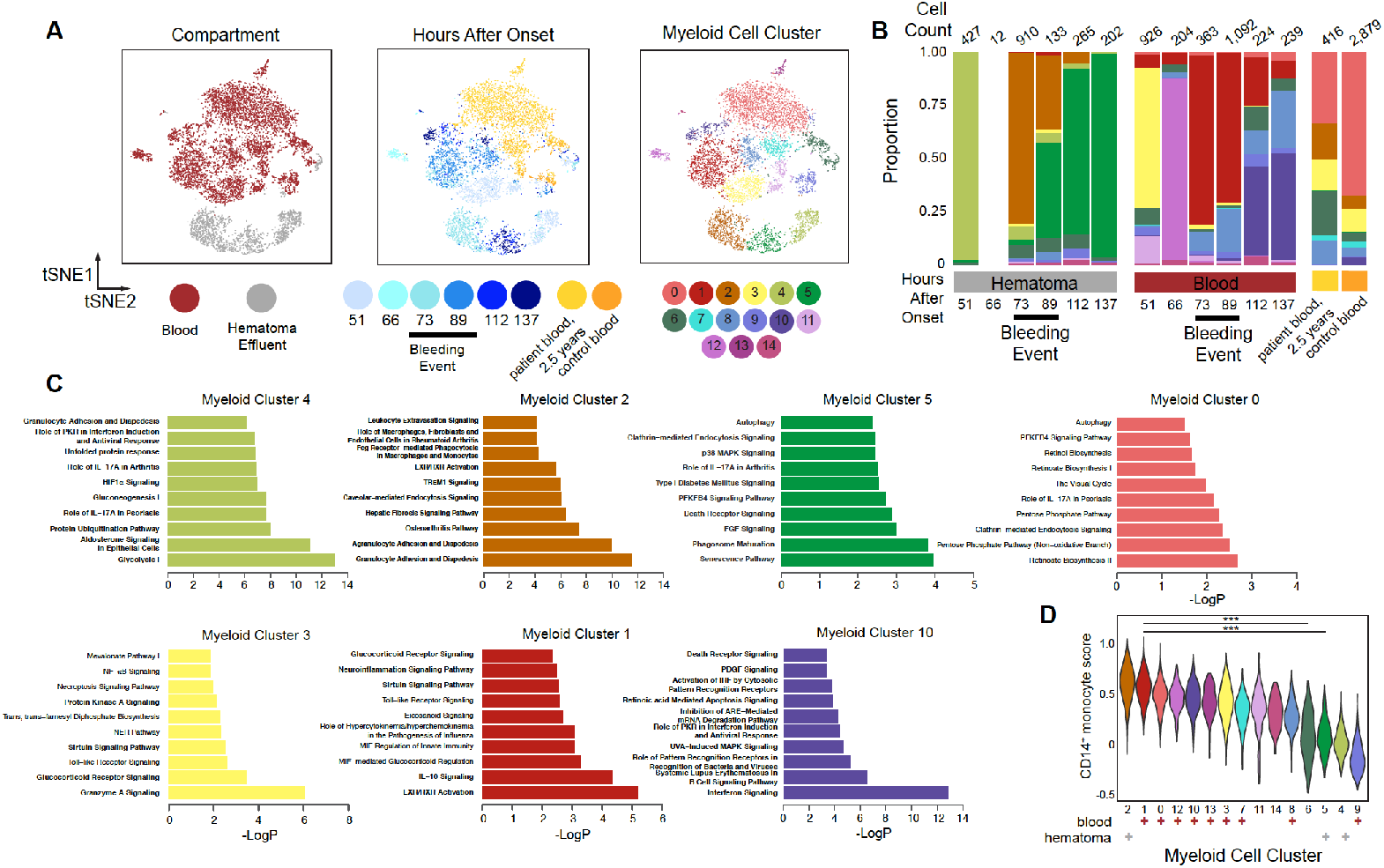
Shifts in prevalence and phenotypes of myeloid cells in hematoma effluent and blood over time. **A**. tSNE plot of re-clustered monocytes, macrophages, and dendritic cells from **Figure 1** (n=8,292 cells). The re-clustered tSNE plot is colored by hematoma or blood (left), time after onset (middle), or by sub-cluster identity. **B**. Stacked frequency plot of new clusters by hours after onset in hematoma and blood. **C**. Top 10 significantly enriched IPA pathways for selected clusters. Myeloid sub-clusters 4, 2, 5 emerge sequentially in hematoma, Myeloid sub-clusters 3, 1, and 10 emerge sequentially in blood, and sub-cluster 0 predominates in patient follow-up at 2.5 years and control blood. Remaining clusters are presented in **Supplemental Figure 8**. Pathways with three or more molecules in the query gene list are bolded. **D**. Violin plot of module scores for an inflammatory monocyte gene signature for each subcluster. All gene modules scored are presented in **Supplemental Figure 9**. Each sub-cluster is annotated as predominantly blood (red +) or hematoma (grey +) in origin below the plot. Annotated adjusted p values were calculated using Wilcoxon rank sum test with Benjamini & Hochberg p value correction, and only select comparisons are annotated on the plot (p_ad_j < 0.001, ***). Full pairwise results for each cluster are shown in **Supplemental Table 7**.

Functional gene enrichment analyses on myeloid sub-cluster defining-genes revealed distinct pathways that were active in each (**Figure 2C,** remaining clusters in **Supplemental Figure 7** and **Supplemental Table 6**). Blood prior to re-bleeding was predominantly comprised of myeloid sub-cluster 3 at 51 hours (enriched for granzyme signaling and glucocorticoid signaling pathways) and myeloid sub-cluster 12 at 66 hours (enriched for iron homeostasis and several T cell interaction pathways). Blood during the re-bleed as detected by catheter drainage (73-89 hours) was predominantly comprised of myeloid sub-cluster 1, enriched for LXR/RXR activation and IL-10 signaling, and then by myeloid subcluster 10, enriched for interferon signaling, at 112 and 137 hours. Sub-clusters unique to control and the patient’s long-term follow-up blood (myeloid subclusters 0 and 13, **Supplemental Figure 6B**) were enriched for autophagy. Overall, sub-clusters predominantly in blood were enriched for a spectrum of pathways involved in innate immune cell functions that changed over time, including TLR signaling, glucocorticoid signaling, EIF2 signaling, and interferon signaling.

Hematoma prior to re-bleeding was predominantly comprised of myeloid sub-cluster 4, which was enriched for glycolysis and HIF-1α signaling. We only recovered 12 myeloid cells at 66 hours in hematoma, limiting our resolution at this time point. There was then a time-dependent shift from dominance by myeloid sub-cluster 2 to myeloid sub-cluster 5. Myeloid sub-cluster 2 was enriched for pathways related to myeloid cell extravasation and tissue infiltration, possibly indicating new infiltration after rebleeding, (**Figure 2C**) and myeloid subcluster 5 was enriched for pathways related to tissue repair and senescence including phagosome maturation, FGF signaling, autophagy, and senescence. This shift is consistent with a transition between dominance of pro-inflammatory activation and recruitment to phagocytosis and perhaps anti-inflammatory functions. Taken together, we found alterations in the overall transcriptional states of myeloid cells in both blood and hematoma, and gene set enrichments suggest these cells, particularly in hematoma, may have differing functions altered by their unique physiological niche that shift over time.

To better contextualize the phenotypes of myeloid sub-clusters, we performed module scoring against a list of curated human monocyte/macrophage gene sets (**Supplemental Table 6, Supplemental Figure 8A**).^11–14^ We performed clustering and principal component analysis (PCA) on these scores to identify modules that explain differences across myeloid clusters. We found that most of these known signatures did not explain differences between myeloid subclusters (**Supplemental Figure 8A**). Nonetheless, we identified four gene signatures that predominantly explain differences between myeloid sub-clusters that were either hematoma or blood in origin, well-describing blood monocytes/macrophages but not those from hematoma (**Supplemental Figure 8B**). Specifically, we found that the predominantly blood sub-clusters (myeloid sub-clusters 0, 1, 3, 6, 7-13) and the sub-cluster that emerged in the hematoma at 73 hours (myeloid sub-cluster 2) scored similarly for a CD14^+^ monocyte gene signature identified recently in blood^12^ (**Figure 2D, Supplemental Figure 9A**). Taken together, our data suggest that myeloid sub-cluster 2 may represent monocytes newly entered into the hematoma due to a recent re-bleed, while other hematoma clusters, like myeloid clusters 4 and 5, are defined by activation states induced by local signals in the hematoma and do not resemble clusters previously defined in human blood or by *in vitro* stimulations.^12–14^ When considered in the context of the timing of the increased draining volume from the catheter and the emergence of a neuron-like cluster at 66hrs, our data may suggest a re-bleeding event prior to the clinically identified changes in hematoma volume detected by a daily CT scan.

Our data also revealed substantial involvement of T cells in the hematoma at all time points. Sub-clustering over all hematoma and peripheral blood T cells from the patient and control blood resulted in 12 T cell sub-clusters that also separated overall by compartment (**Figure 3A, Supplemental Figure 9**). As with monocytes, we found that T cell frequencies were dynamic over the course of ICH in hematoma (**Supplemental Table 7**). In the blood, there was an initial shift in the dominant cluster from T cell sub-cluster 3 to T cell sub-cluster 2 prior to the re-bleed, which then remained highly frequent for the remaining time points (**Figure 3B**). Several sub-clusters were also predominantly blood in origin (subclusters 0, 2, 4, 10, and 11). T cell sub-clusters 2 and 3, which were enriched for IL-17A and GADD45 signaling, respectively, also emerged in the hematoma at 73 hours. The emergence of these clusters in hematoma at this time paralleled the emergence of a more blood-like monocyte sub-cluster at the same time point, further supporting evidence this was the beginning of the rebleeding.

**Figure 3.**
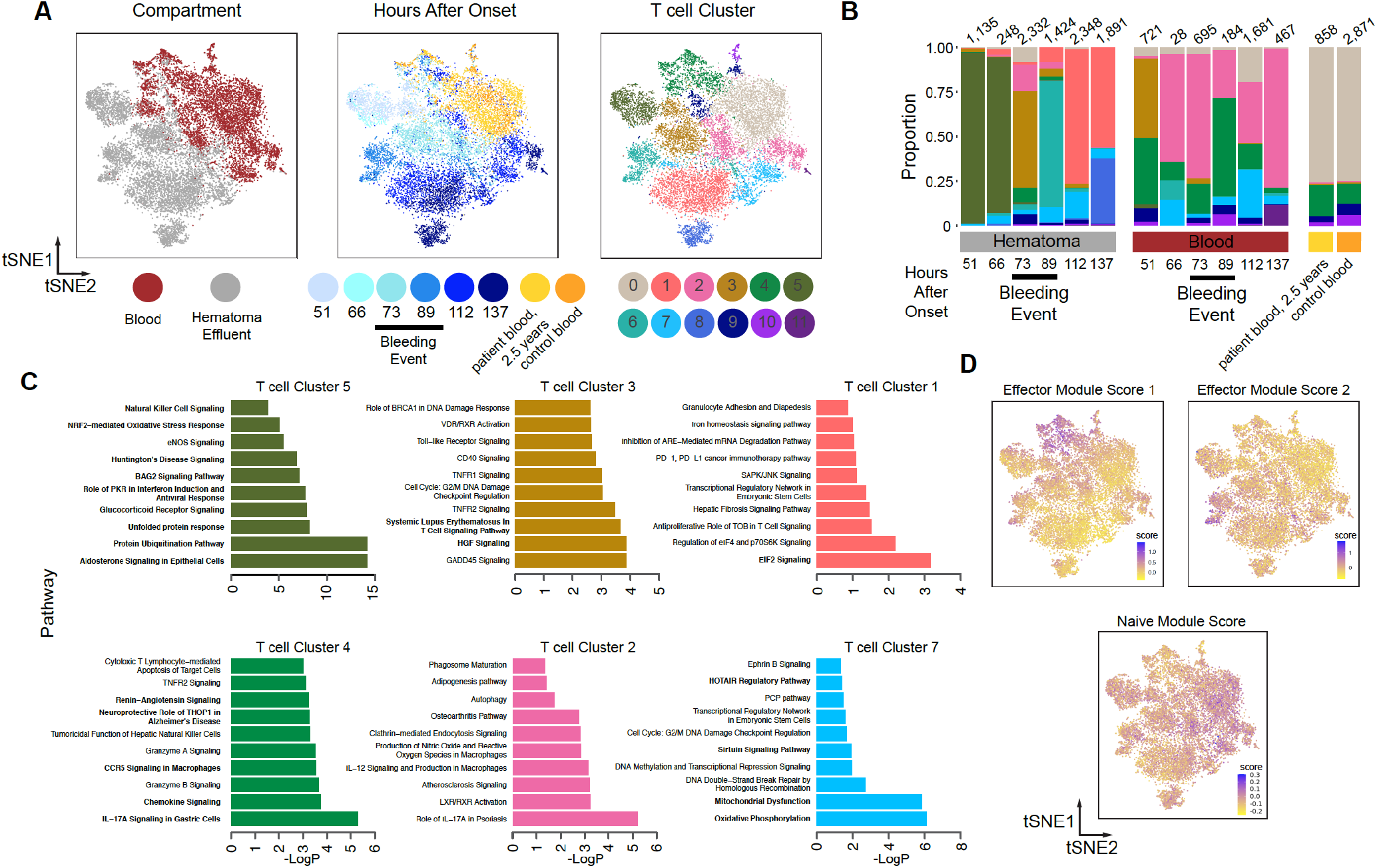
Shifts in prevalence and phenotypes of T cells in hematoma effluent and blood over time. **A**. tSNE plot showing re-clustered T cells (n=16,883 cells) from **Figure 1**. The re-clustered tSNE plot is colored by hematoma or blood (left), time after onset (middle), or T cell sub-cluster identity (right) determined by re-clustering analysis. **B**. Stacked frequency plot of new T cell clusters by hours after onset in blood and hematoma. **C**. Top 10 significantly enriched IPA pathways for selected clusters. T cell sub-clusters 5, 3, and 1 emerge in hematoma and T cell sub-clusters 4, 2, and 7 are found in blood. Remaining clusters are presented in **Supplemental Figure 11**. Sub-cluster 0 was defined by two marker genes (TXNIP and LTB). Pathways with three or more molecules in the query gene list are bolded. **D.** tSNE plots colored by each indicated module score.

Several sub-clusters of T cells were predominantly found in hematoma (**Supplemental Figure 9B**) (e.g., T cell sub-clusters 1, 3, 5, 6, and 8). T cell subcluster 5, prevalent in early hematoma samples, was enriched for heat shock proteins, protein ubiquitination, and eNOS signaling (**Figure 3D, Supplemental Table 8**). T cell sub-cluster 3, which emerged at 73 hours, was enriched in GADD45 signaling and several immune activation pathways (**Figure 3C,** additional clusters in **Supplemental Figure 10**). T cell sub-cluster 1, emerging at later time points in hematoma, shows expression of genes involved in transcriptional regulation, anti-proliferation, and iron homeostasis. Interestingly, we did not find clear patterns of chemokine and receptor expression^15^ in specific clusters in either myeloid or T cell sub-clusters; we did find, however, elevated expression of CXCR4 consistently across myeloid and T cell sub-clusters (**Supplemental Figure 11)**. Overall, we found that T cells that originated in blood are enriched for innate immune interactions, IL-17 signaling, chemokine signaling and sirtuin signaling, whereas those predominantly from hematoma are enriched for stress response pathways, diapedesis and neuroinflammation. Both the patient’s long-term follow up blood sample and the healthy control blood samples were predominated by T cell sub-cluster 0, which was defined by two marker genes (TXNIP and LTB).

To better contextualize our T cell sub-clustering results, we again performed module-scoring analysis with known T cell gene signatures previously derived from human transcriptional data (**Supplemental Table 10**). Unsupervised clustering followed by PCA on module scores revealed several modules that describe variation across T cell sub-clusters (**Supplemental Figure 12A**). We found that three gene modules that describe effector functions in T cells drove major axes of variation in our dataset (**Figure 3D**). One signature is associated predominantly with T cell sub-cluster 4 and is defined by T cell effector genes like GNLY, CCL4, and GZMK (Effector Module Score 1, **Supplemental Figure 12B**). The other is associated with T cell sub-cluster 6, which is a hematoma cluster emerging during rebleeding, and is defined by T cell effector genes like CXCR4, TXN, and CSTB (Effector Module Score 2, Figure 3d, **Supplemental Figure 12B**). Generally, cells in clusters on the right in the tSNE plot shown in **Figure 3D**, including T cell sub-clusters 0, 2, and 7 (found predominantly in blood), score higher for a naïve T cell module. Overall, these data – combined with our pathway enrichment results – suggest that T cells in the predominantly hematoma-derived clusters may be enriched for unique T cell functions, including several related to effector functions and stress responses.

Based on the emergence of several blood-like monocyte and T cell subclusters in the hematoma at 73 hours, as well as increased drainage volume and cellularity, we estimate that the time of re-bleeding was between 66 and 73 hours after ICH onset, a full day prior to detection by CT scan. It is also intriguing that neurons emerged in the hematoma effluent immediately prior to this event, but the consequences of this is unknown as the patient continued to improve. Monitoring of specific peripheral blood biomarkers found in effluent could thus potentially enable earlier detection of a re-bleeding event to guide a more effective treatment paradigm. We caution, however, that our study presents only one patient’s ICH trajectory; nevertheless, it suggests that larger scale scRNA-seq studies could provide more insight into specific blood cell states that may better inform patient treatment and outcome and highlight pathways for future research.

Our transcriptional analyses of leukocytes over the course of ICH in this patient suggest coordinated, dynamic stages of monocyte and T cell responses within the hematoma that were altered after asymptomatic re-bleed, indicating rapid adaptation and response to local changes in the hematoma tissue milieu. Additionally, we studied the cellular composition of the blood of this patient more than two years after ICH, and our data suggest that the peripheral blood of this patient, who had an excellent outcome, is comparable to that of age-matched controls.

The dynamics of leukocyte infiltration and activation after ICH have been difficult to study in patients, especially at the site of injury.^16^ At present, our understanding is derived primarily from rodent models and peripheral measures; how the kinetics in these models compares to those in humans is unknown.^17,18^ Using the infrastructure of the MISTIE III trial, we characterized the local cellular response to ICH and an associated re-bleeding event in the brain of a living patient at single-cell resolution for the first time. We characterized thousands of single cells from both blood and hematoma effluent at several points after ICH onset, allowing us to identify cellular phenotypes that were dynamic in both the T cell and myeloid lineages. Most notably, the emergence of myeloid sub-cluster 2 and T cell sub-cluster 2 in the hematoma after re-bleeding, followed by the sequential emergence of new cell states, suggests that newly infiltrating immune cells adapt transcriptionally to dynamic injury events in the brain.

Crucially, our work demonstrates how rapidly immune cell activation states change after brain injury in humans and sets the stage for framing critical time periods for future studies. Our data, while we caution is a small case study, provide proof-of-principle that, in the near future as personalized medicine evolves, this and other approaches could be used clinically to interrogate cellular responses and uncover meaningful clinical events in patients to help guide treatment decisions.

## METHODS

The methods are described in the Supplemental Materials.

### Study approval

The patient was enrolled in the MISTIEIII trial (NCT01827046) under an IRB approved protocol including sample collection. Control donors (n=4, 2 males and 2 females, ages 63-93) were recruited and samples were also collected under an IRB approved protocol.

## AUTHOR CONTRIBUTIONS

LHS designed the study in consultation with DFH, JCL, BAG, MHA and AKS. LHS, AKS, and JCL supervised the study at all stages. Sample processing was performed by MHA and HEB. BAG, MHA, JHD, NHP performed Seq-Well and associated sequencing with assistance from IF, EM and RSD. BAG performed transcriptional analysis with input from LHS, MHA, JCL and AKS. MHA performed mass cytometry and mass cytometry analysis. DFH, IAA, and MZ led the MISTIE III trial internationally and provided trial data for this study and CCM and LHS led trial at Yale and provided patient samples and additional data. BAG, MHA, JCL, AKS, and LHS wrote the manuscript with critical input from all authors.

## ACKNOWLEDGEMENTS

MISTIEIII (U01NS080824, PI Hanley), Trial innovation Network (U01NS080824 PI Hanley) and JHU CTSA (U24TR001609 Hanley section PI), ICHseq (R01NS097728, PI Sansing), NRSA postdoctoral fellowship (F32-AI136459, Goods), AHA postdoctoral fellowship (17POST33660872, Askenase), Swanson Biotechnology Center at the Koch, Searle Scholars Program (Shalek), the Beckman Young Investigator Program (Shalek), a Sloan Fellowship in Chemistry (Shalek), and NIH 5U24AI118672 (Shalek).

## Supplemental Appendix

### ICHSeq Investigators

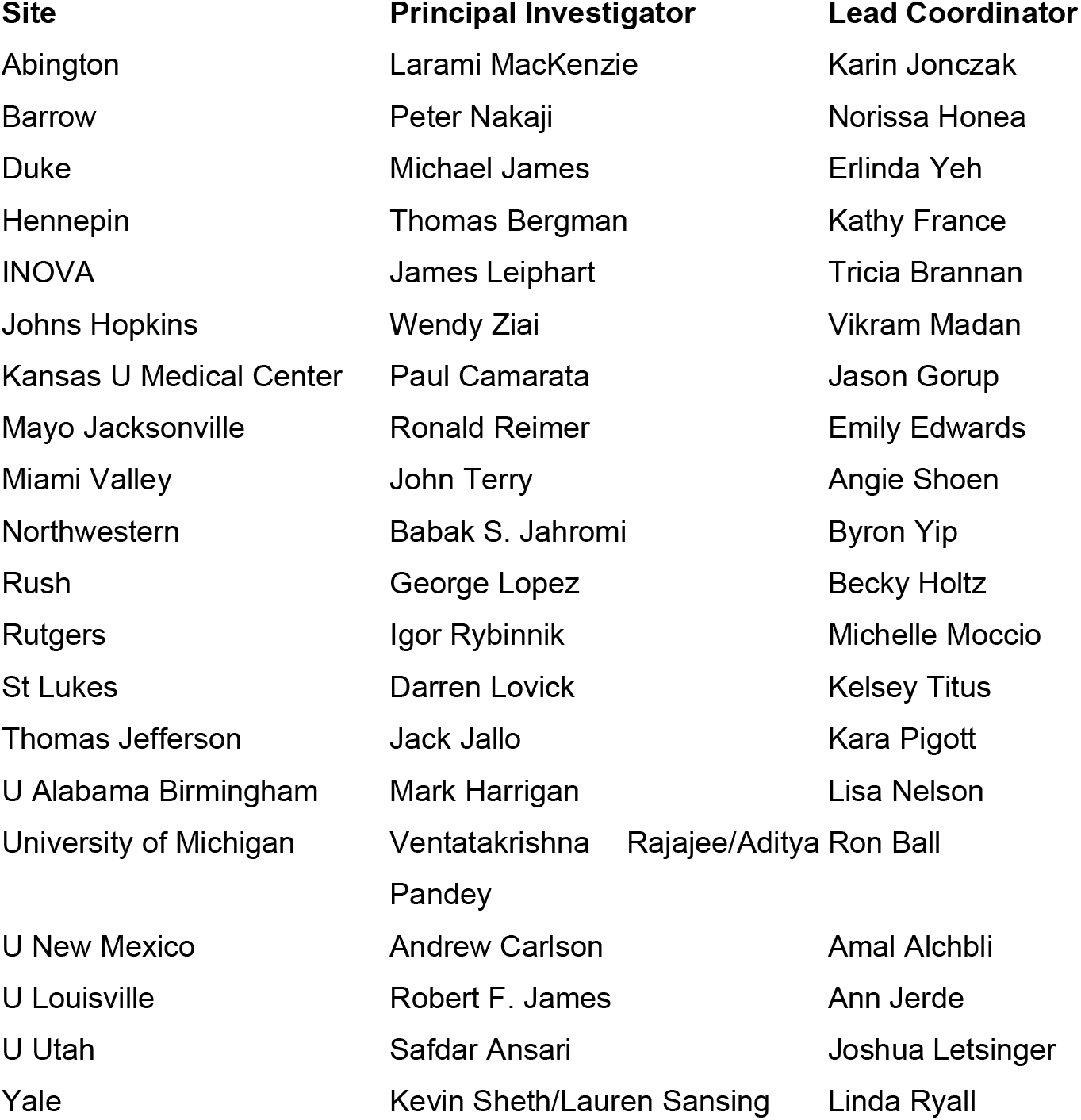

### BIOS and MTI:M3 Investigators

|  |  |
| --- | --- |
| Johns Hopkins | Paul Nyquist |
| Mass General | Josh Goldstein |
| Johns Hopkins | Vikram Madam |
| Johns Hopkins | Christina Grabartis |
| Johns Hopkins | Tracey Economas |
| Johns Hopkins | Wendy Ziai |
| Manchester | Adrian R. Parry-Jones |

### Supplemental Methods

#### Patient enrollment

The patient was enrolled in the MISTIEIII trial (NCT01827046) under an IRB approved protocol including sample collection. Control donors (n=4, 2 males and 2 females, ages 63-93) were recruited and samples were also collected under a IRB approved protocol.

#### Patient hematoma effluent and blood collection and processing

Each sample was centrifuged at *500g* for 10 minutes and the supernatant was removed. Then each sample was resuspended in HBSS (Gibco) and treated with 2.5 Units/ml of Benzonase (Sigma #E1014) for 10 minutes at room temperature. Samples were then passed through a 70-micron filter to remove tissue debris, washed with 25 ml of HBSS, and centrifuged at 300*g* for 10 minutes. After supernatant was removed, platelets and most erythrocytes were removed from these samples using a LeukoLock filter (Life Technologies) as previously described. Briefly, after the sample passed through the filter binding leukocytes, the filter was backflushed to recapture the leukocytes. This backflush was centrifuged at 300g for 8 minutes and supernatants were removed until the mixed leukocyte/erythrocyte cell pellet composed approximately 50% of the total volume of the sample. Granulocytes were removed from this sample using the RosetteSep Human Granulocyte Depletion Cocktail (Stem Cell Technologies) and subsequent Ficoll density gradient centrifugation according to the manufacturer’s protocol. Finally, the resulting cell pellets, which still contained trace numbers of erythrocytes, were subjected to two rounds of erythrocyte lysis at room temperature for 5 minutes each using BD PharmLyse buffer, washed, and resuspended in RPMI containing 10% FBS.

#### Generation of single-cell RNA-sequencing (scRNA-seq) data with Seq-Well

Seq-Well was performed as described previously.^1^ About 15,000 cells were loaded onto each array preloaded with uniquely-barcoded mRNA capture beads (ChemGenes). Arrays were washed with protein-free RPMI media, then sealed with polycarbonate membranes. Arrays were incubated at 37°C for 30 minutes to allow membranes to seal, then transferred through a series of buffer exchanges to allow for cell lysis, transcript hybridization, bead washing, and bead recovery from arrays post membrane removal. Reverse transcription was performed with Maxima H Minus Reverse Transcriptase (ThermoFisher), excess primers were removed using an Exonuclease I digestion (New England Biolabs), and whole transcriptome amplification (WTA) by PCR was performed using KAPA Hifi PCR Mastermix (Kapa Biosystems). WTA product was purified using Agencourt Ampure beads (Beckman Coulter) and 3’ digital gene expression (DGE) sequencing libraries were prepared using Nextera XT (Illumina). Two arrays worth of sequencing libraries were sequenced per NextSeq500/550 run using a 75 cycle v2 sequencing kit (Illumina) with a paired end read structure (R1: 20 bases; I: 8 bases; R2: 50 bases) and custom sequencing primers.

#### scRNA-seq computational analyses

Raw sequencing data was demultiplexed and aligned to the Hg19 genome using publicly available scripts on Terra (scCloud/dropseq_workflow, version 11, https://app.terra.bio/). The resulting UMI-collapsed digital gene expression matrices were used as input to Seurat (v3) for further analyses in R (v3.6.2). Initial clustering was performed, and on the basis of marker genes identified using Seurat’s Wilcoxon ranksum test, red blood cells were identified by expression of hemoglobin genes and excluded from further analysis, resulting in a total of 31,722 cells across all conditions. All data were normalized, the top 2,000 variable genes identified, and the data was clustered at a resolution of 2. Marker genes for each cluster were identified using Seurat’s Wilcoxon rank-sum test, and broad immune cell types were labeled by comparing to known marker genes that have been previously published, and clusters were collapsed into labels as shown in Figure 1.^2,3^

For sub-clustering analyses, T cells or myeloid cells were analyzed separately. Each cell-type was re-normalized, the top 2,000 variable genes were identified, and the data was clustered across several resolutions to identify resolutions that produced non-redundant clusters (resolution = 0.6 for T cell sub-clustering, and resolution = 0.6 for myeloid sub-clustering) as determined by marker gene identification using Seurat’s Wilcoxon rank-sum test. Notebooks to reproduce all analyses performed in R will be available for download (https://github.com/ShalekLab).

Functional enrichment was performed using Ingenuity Pathway Analysis (Qiagen, Version 47547484) using adjusted p-values of significant genes in each cluster as input, and significant canonical pathways were reported if their p-value was significant (p_adj_>0.05). If three or more molecules of the pathways were identified in the input gene lists, these pathways are bolded in the main manuscript figures. For creating gene module scores, gene lists were manually curated from existing literature sources (**Supplemental Table 6 and 10**) and scores were generated in R using Seurat.

#### Mass cytometry sample preparation

Cell suspensions from each time point was stained for mass cytometry and fixed immediately upon isolation.^4^ Antibody mixes (**Supplemental Table 2**) were prepared fresh each day in BSA stain buffer (BD, #554657) and centrifuged through a 0.1 micron filter (Millipore #UFC30VV00) at 12,000*g* for 4 minutes; this process reduced nonspecific labeling of cells with free isotopes. Up to 3 x 10^6^ cells in RPMI containing 10% FBS were transferred to a 1.5 ml Eppendorf tube and washed with phosphate-buffered saline (PBS). Cell suspensions were centrifuged at 300g for 8 minutes and supernatant was removed from the cell pellet. For all centrifugation steps, a fixed rotor microcentrifuge was used and supernatants were carefully removed by pipette. To mark cells for viability, cell pellets were resuspended in 1 μM Cell-ID cisplatin (Fluidigm #201194) and incubated for 3 minutes at room temperature. Cells were washed with 1 mL of BSA stain buffer, centrifuged at *300g* for 8 minutes, and supernatant was carefully removed. Cell pellets were resuspended in antibody mix and incubated on ice for 40 minutes. Cell pellets were washed with 1 mL of BSA stain buffer, centrifuged, resuspended in Fix/Perm buffer from the FoxP3 staining kit (eBiosciences, #00-5523-00), and incubated overnight at 4°C. The next morning, samples were centrifuged at 800*g* for 5 minutes, resuspended in 300 μL BSA stain buffer, and stored at 4°C until all timepoints had been collected. Then, all samples were centrifuged and resuspended in 125 nM Cell-ID Intercalator-IR (Fluidigm #201192A) diluted in Fix/Perm buffer, and incubated overnight at 4°C. Immediately before running each sample on the mass cytometer, that sample was centrifuged, supernatant was removed, and the pellet was resuspended in deionized water.

#### Data analysis for mass cytometry

Initial gating was performed to identify CD45^hi^ cisplatin^lo^ viable leukocytes, after first using Iridium intercalator to exclude doublets and non-cellular debris. Cells meeting these criteria were then normalized using bead standards to account for variations in cytometer sensitivity as previously described.^4^ Cell populations were identified using FlowJo software (v10, Treestar) according to the following marker combinations for each: CD4 T cells (CD3^+^CD19^−^CD4^+^), CD8 T cells (CD3^+^CD19^−^CD8a^+^), NK cells (CD3^−^ CD56^+^CD11b^lo^), B cells (CD3^−^CD56^−^CD11b^−^CD19^+^), monocytes (CD3^−^CD56^−^ CD11b^hi^CD19^−^CD66a^−^CD14^+^CD11c^+^), neutrophils (CD3^−^CD56^−^CD11b^hi^CD19^−^ CD66a^+^HLA-DR^lo^).

#### Bulk RNA-sequencing sample preparation and analysis

Up to 5 x 10^6^ leukocytes isolated from each sample by the above method were used for downstream sorting. Cells were first lightly fixed in 1 mL of Cell Cover, a light preservative that maintains RNA integrity (Anacyte #800-250), on ice for 10 minutes, then centrifuged. CD3^+^ cells were separated from total cell suspensions by magnetic selection (StemCell, #17851) according to manufacturer’s instructions. CD3^+^ cells were stained for flow cytometry using CD45 (Tonbo, #35-0459-T100), CD25 (BD Biosciences, #555432), CD8 (Tonbo, #60-0088-T100), CD127 (Biolegend, #351316), CD2 (BD Biosciences, #562638), CD4 (Tonbo, #75-0049-T100), CD11b (Tonbo 30-0112-U500), CD20 (Biolegend, #302349), CD56 (Biolegend, #318320), and viability dye (Life Technologies, #L34972). CD45^hi^ CD11b^−^ CD56^−^ CD20^−^ CD2^+^ CD4^+^ CD8^−^, CD127^hi^ CD25^−^ live cells were sorted for CD4 T cell library construction. CD3^−^ cell fractions were stained for flow cytometry with CD45 (Tonbo Biosciences, #50-0459-T100), CD11b (Tonbo, #35-0118-T100), CD14 (Biolegend, #301820), CD16 (BD Biosciences, #560474), CD66a/c/e (Biolegend, #342310), CD2, (Biolegend, #300204), CD20 (Biolegend, #302349), CD56 (Biolegend, #318320), and viability dye. CD45^hi^ CD2^−^ CD56^−^ CD20^−^ CD66^−^ CD11b^hi^ CD14^hi^ CD16^lo^ cells were sorted for monocyte library construction. Monocyte and CD4 populations were sorted using a FACSaria II (**Supplemental Figure 7A and B**) directly into RNA lysis buffer composed of 200 μL RA1 buffer (Macherey-Nagel, #RA1) freshly spiked with 2% TCEP (Thermofisher, #77720).

RNA-sequencing libraries were generated as previously described.^5^ Briefly, RNA was extracted using the NucleoSpin RNA XS Kit (Macherey-Nagel, #740902) according to the manufacturer’s instructions. Smart-Seq2 cDNA synthesis was performed with the following modifications: 1) input RNA was normalized prior to cDNA generation by diluting to ~1,000 cells per reaction, 2) reverse transcription was performed with Superscript III (ThermoFisher, #18080-085) in place of Superscript II according to the manufacturer instructions. Paired-end sequencing libraries were prepared using the Nextera XT DNA sample Prep Kit (Illumina, #FC-131) according to the manufacturer’s instructions. Libraries were pooled in an equimolar ratio and sequenced on a NextSeq500/550 sequencer (Illumina) using a 75 cycle v2 sequencing kit with a paired end read structure. Following sequencing, BAM files were converted to merged, demultiplexed FASTQs. Paired-end reads were mapped to the UCSC hg19 genome using STAR and RSEM. SSGSEA was performed on the Hallmark gene sets using the “SSGSEAProjection” module version 9.1.1 on the GenePattern website (https://www.genepattern.org/) with the following parameters: weighting exponent = 0.75, minimum gene set size = 10, and combine mode = combine.add. Heatmaps of resulting ssGSEA results were generated using Morpheus.

#### Flow cytometry analysis of blood and hematoma leukocyte populations

Cell suspensions were generated as described above, except granulocytes were not depleted prior to red blood cell lysis. 10^6^ whole blood leukocytes from the resulting cell suspensions were stained for flow cytometry with CD14 (eBioscience, 11-0149-42), CD25 (BD Biosciences, 555432), CD66a/c/e (Biolegend, 342310), CD45 (Biolegend, 304012), CD3 (BD Biosciences, 560176), CD4 (Tonbo, 75-0049-T100), CD8 (BD Biosciences, 560774), CD11b (Biolegend, 301336), and viability dye (Life Technologies, L34972). Flow cytometry was performed on a BD Fortessa (**Supplemental Figure 5A**). After eliminating dead cells using viability dye, the following populations were identified using FlowJo software (Treestar): CD4 T cells (SSC^lo^, CD45^+^, CD66^−^, CD3^+^, CD11b^−^, CD4^+^, CD8^−^), CD8 T cells (SSC^lo^, CD45^+^, CD66^−^, CD3^+^, CD11b^−^, CD8^+^, CD4^−^), CD14^hi^ monocytes (SSC^lo^, CD45^+^, CD66^−^, CD11b^+^, CD3^−^, CD14^hi^, CD16^lo^), CD16^hi^ monocytes (SSC^lo^, CD45^+^, CD66^−^, CD11b^+^, CD3”, CD16^hi^, CD14^lo^), and neutrophils (SSC^hi^, CD45^lo^, CD66^+^, CD16^+^).

#### Statistical analysis

Statistical analyses for comparing cell frequencies obtained with CyTOF or Seq-Well were performed using Prism (8.4.3). P values were calculated by repeated measures analysis of variance (ANOVA) followed by post hoc test (Kruskal-Wallis, nonparametric) and also where indicated standard error bars are shown. For comparison of module scores, analyses were performed in R (v3.6.2). Module scores between myeloid sub-clusters were compared by Kruskal-Wallis rank sum test followed by pairwise comparisons using Wilcoxon rank sum test with Benjamini & Hochberg p value correction.

#### Data availability

Raw sequencing data will be made available through Gene Expression Omnibus (Study ID: #####) concurrent with publication, as well as through The Alexandria Project (https://singlecell.broadinstitute.org/single_cell?scpbr=the-alexandria-project).

### Supplemental Figures

**Supplemental Figure 1.**
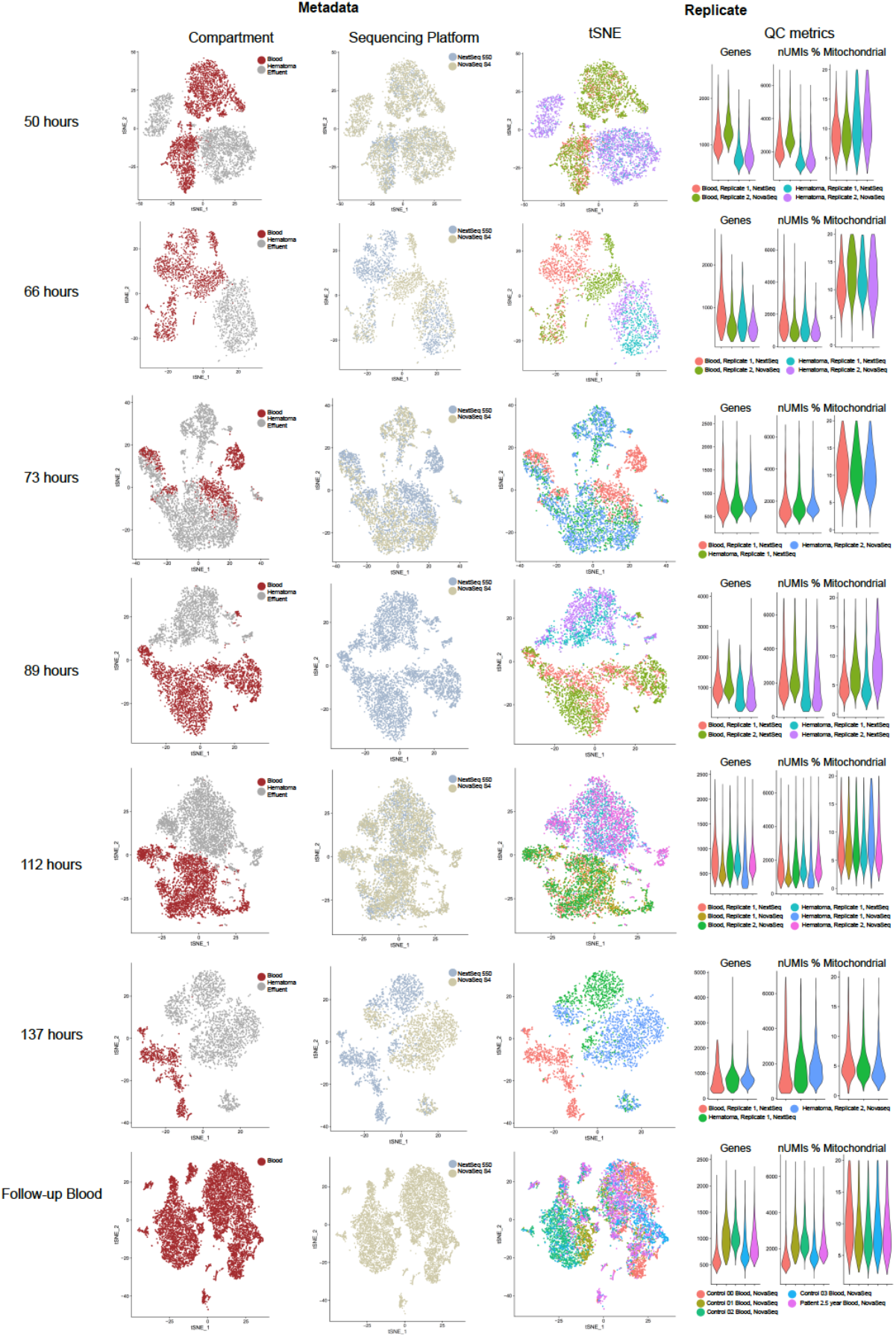
Reproducibility across replicates. tSNEs of data for each time point post ICH are shown and colored by metadata (compartment or sequencing platform) and replicatearray. Violin plots of quality control metrics including number of unique molecular identifiers (nUMIs), number of genes detected (Genes), and fraction of mitochondrial genes for each SeqWell array.

**Supplemental Figure 2.**
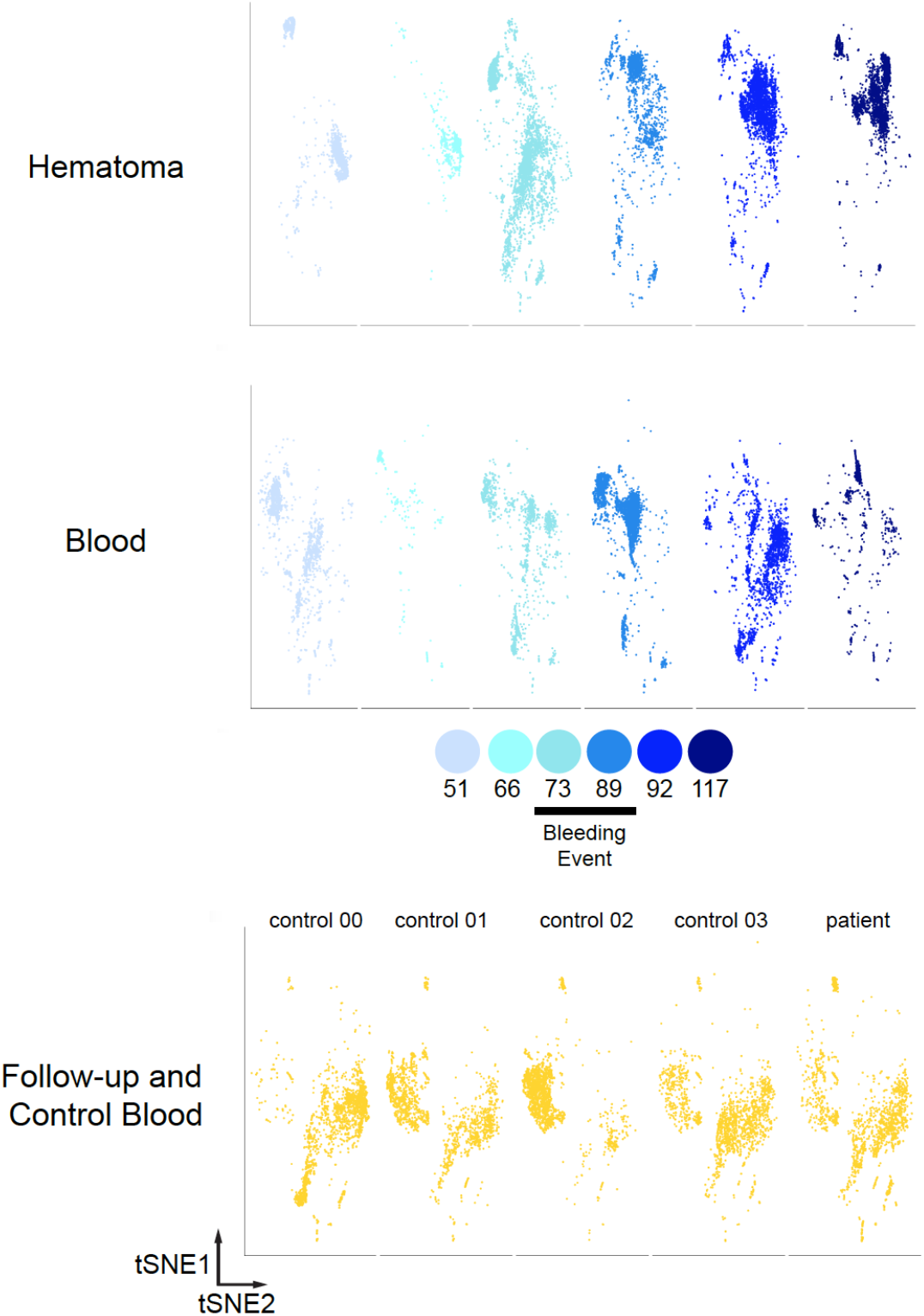
tSNE of all data shown by hematoma (top), blood (middle), or control and patient 2.5 year follow-up. tSNEs are also shown by individual time point, showing cell yield for each array.

**Supplemental Figure 3.**
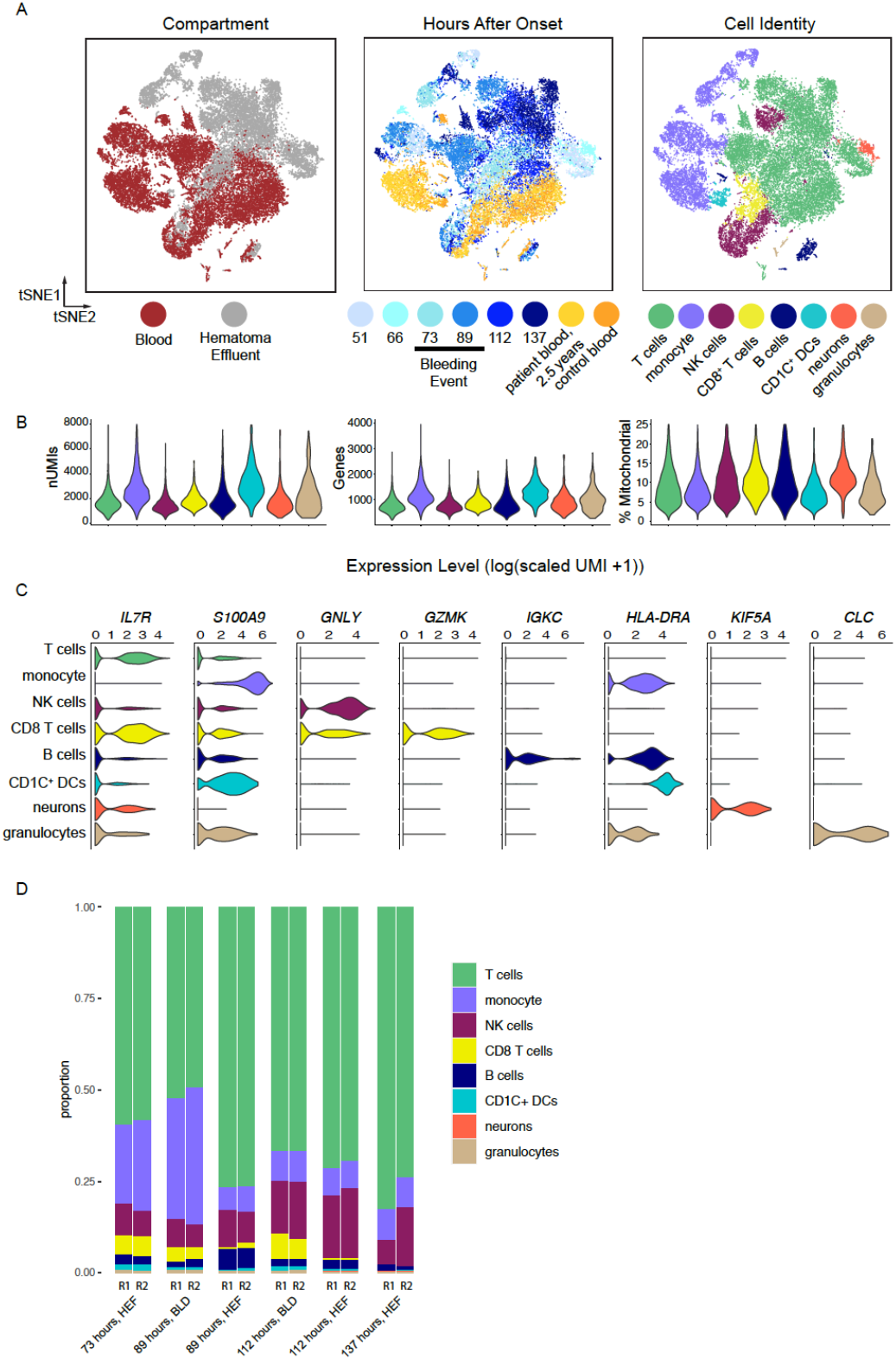
tSNE of all data clustered including all cell clusters with key cell type identification genes as feature plots. **A**. tSNE colored by compartment, time, and cell type reproduced from Figure 1. **B**. Violin plots of quality control metrics including number of unique molecular identifiers (nUMIs), number of genes detected (Genes), and fraction of mitochondrial genes per cell plotted by cell type. **C**. Violin plots show increased expression of selected marker genes for each cluster. **D.** Frequency of each cell type is shown for replicate arrays.

**Supplemental Figure 4.**
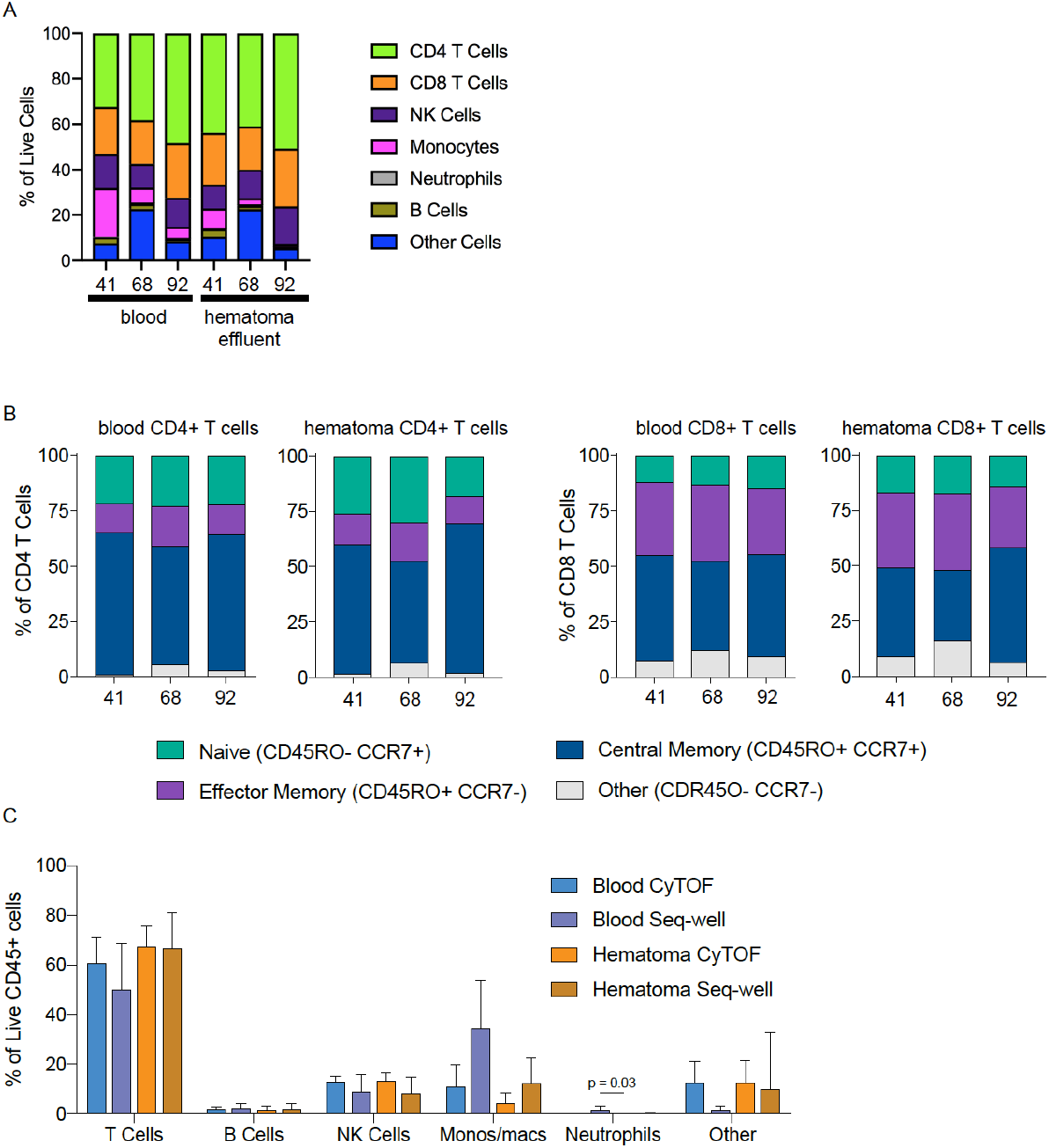
Mass cytometry analysis of hematoma and blood. **A**. Stacked bar chart of percent of live cells. Cell populations were identified according to the following marker combinations for each: CD4 T cells (CD3^+^CD19^−^CD4^+^), CD8 T cells (CD3^+^CD19^−^CD8a^+^), NK cells (CD3^−^ CD56^+^CD11b^lo^), B cells (CD3^−^CD56^−^CD11b^−^ CD19^+^), monocytes (CD3^−^CD56^−^ CD11b^hi^CD19^−^CD66a^−^CD14^+^CD11c^+^), neutrophils (CD3^−^CD56^−^CD11b^hi^CD19^−^ CD66a^+^HLA-DR^lo^). **B.** Stacked bar charts of indicated T cell subsets. **C**. Comparison of cell type frequencies between indicated platform (CyTOF or Seq-Well).

**Supplemental Figure 5.**
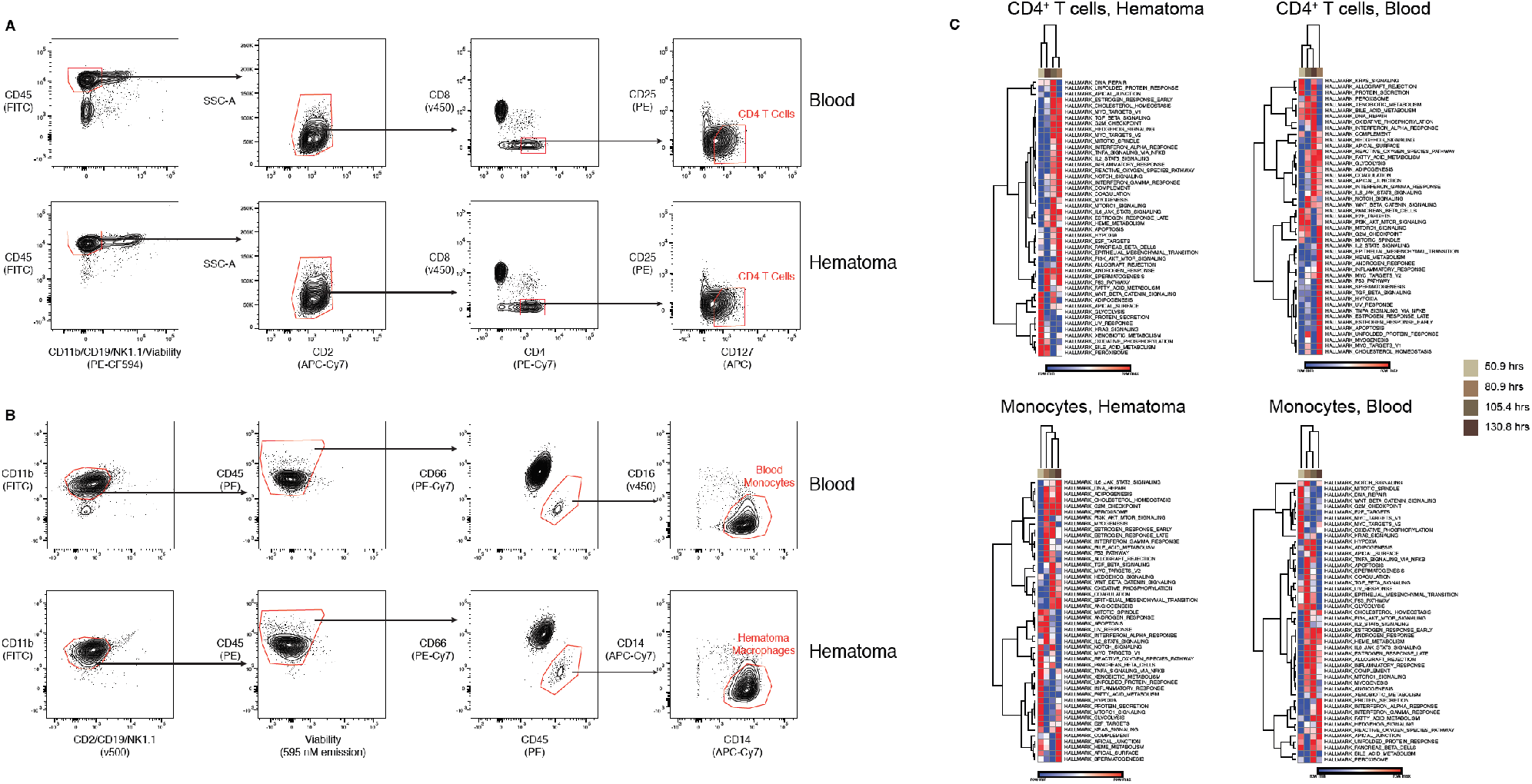
Patient bulk RNA-seq. **A.** Representative sorting strategy for CD4^+^ T cells isolated from blood and hematoma for bulk RNA-seq. **B.** Representative sorting strategy for monocytes isolated from blood and hematoma for bulk RNA-seq. **C.** Heatmaps of SSGSEA results from bulk RNA-sequencing on monocytes and CD4^+^ T cells from blood or hematoma for non-overlapping time points from this patient.

**Supplemental Figure 6.**
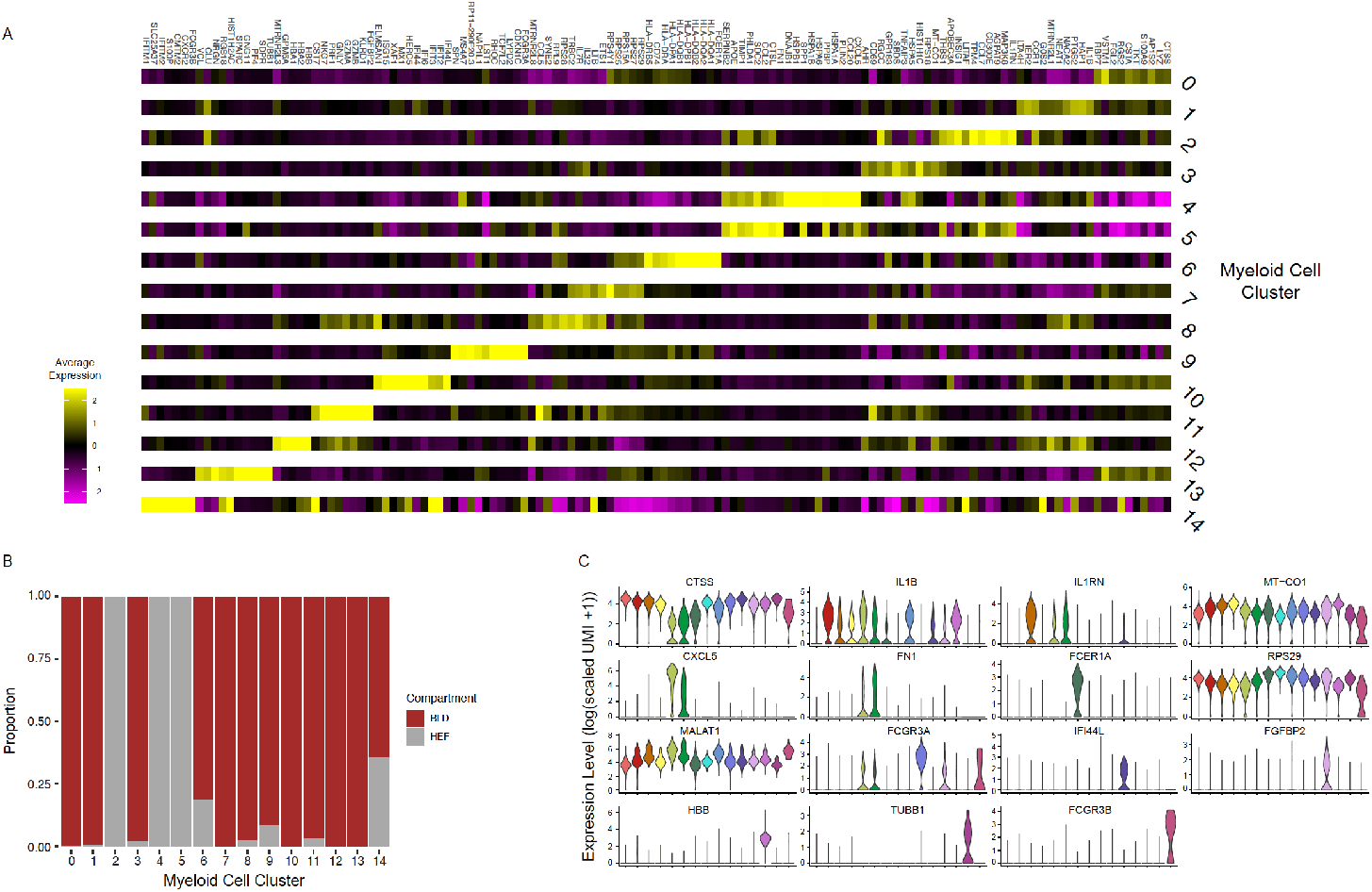
Myeloid re-clustering analysis. **A**. Heatmap of top ten differentially expressed genes for each myeloid sub-cluster. **B**. Stacked bar chart showing frequency of compartment (blood or hematoma) for each myeloid subcluster. **C**. Violin plots of selected marker genes for each myeloid sub-cluster.

**Supplemental Figure 7.**
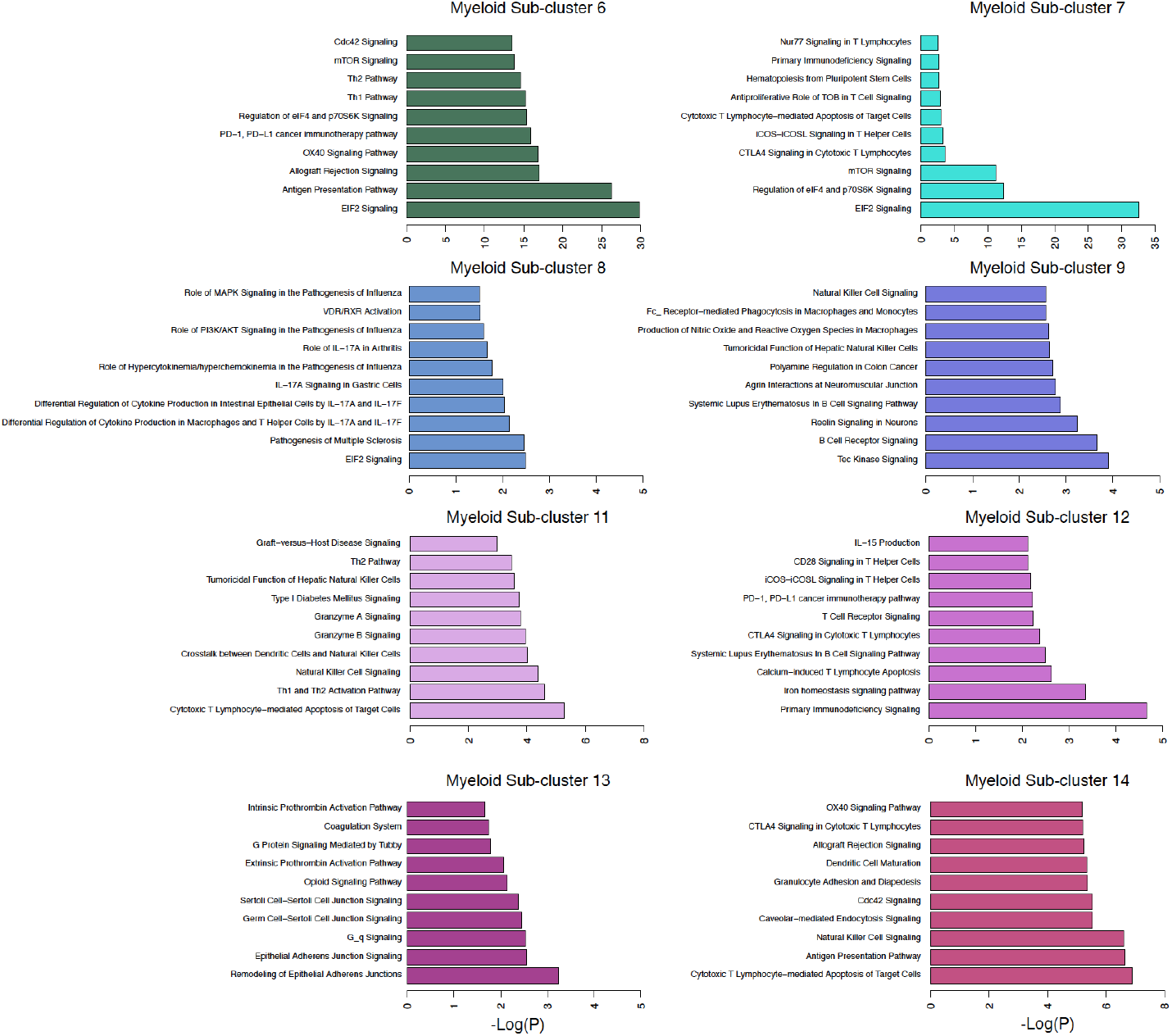
Waterfall plots of top 10 enriched pathways by IPA for remaining new myeloid clusters. Results, including genes in each pathway, are also shown in Table S5.

**Supplemental Figure 8.**
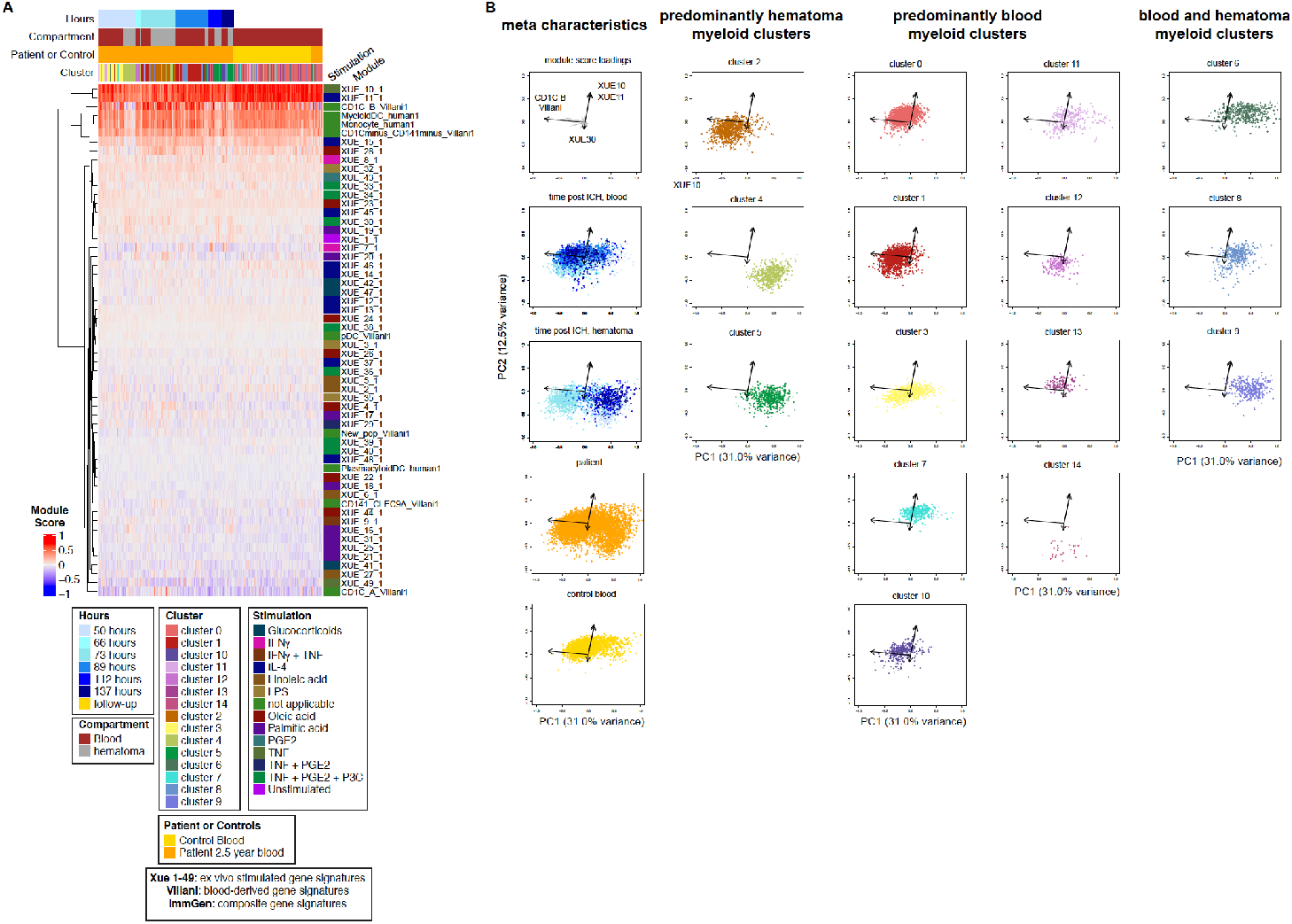
**A**. Gene expression module scores were calculated for selected gene modules from the literature for each cell in our myeloid data and clustered hierarchically. The heatmap is annotated by compartment (blood or hematoma), time, source (control or patient), and myeloid sub-cluster. Each module scored over is annotated by primary stimulation condition. B. Principal component analysis was performed on all module scores. Data are shown across the first two principal components and all module variable loadings are projected onto the top left plot and each subsequent plot shows only the top four module loading (black arrows). Data from each myeloid sub-cluster is visualized separately (sub-clusters 0-14), or as a function of time in either blood or hematoma, and as patient derived or control blood derived. Plots are grouped by metadata characteristics, predominantly blood, predominantly hematoma, or both. Gene signatures used for module scoring are described in **Table S6**.

**Supplemental Figure 9.**
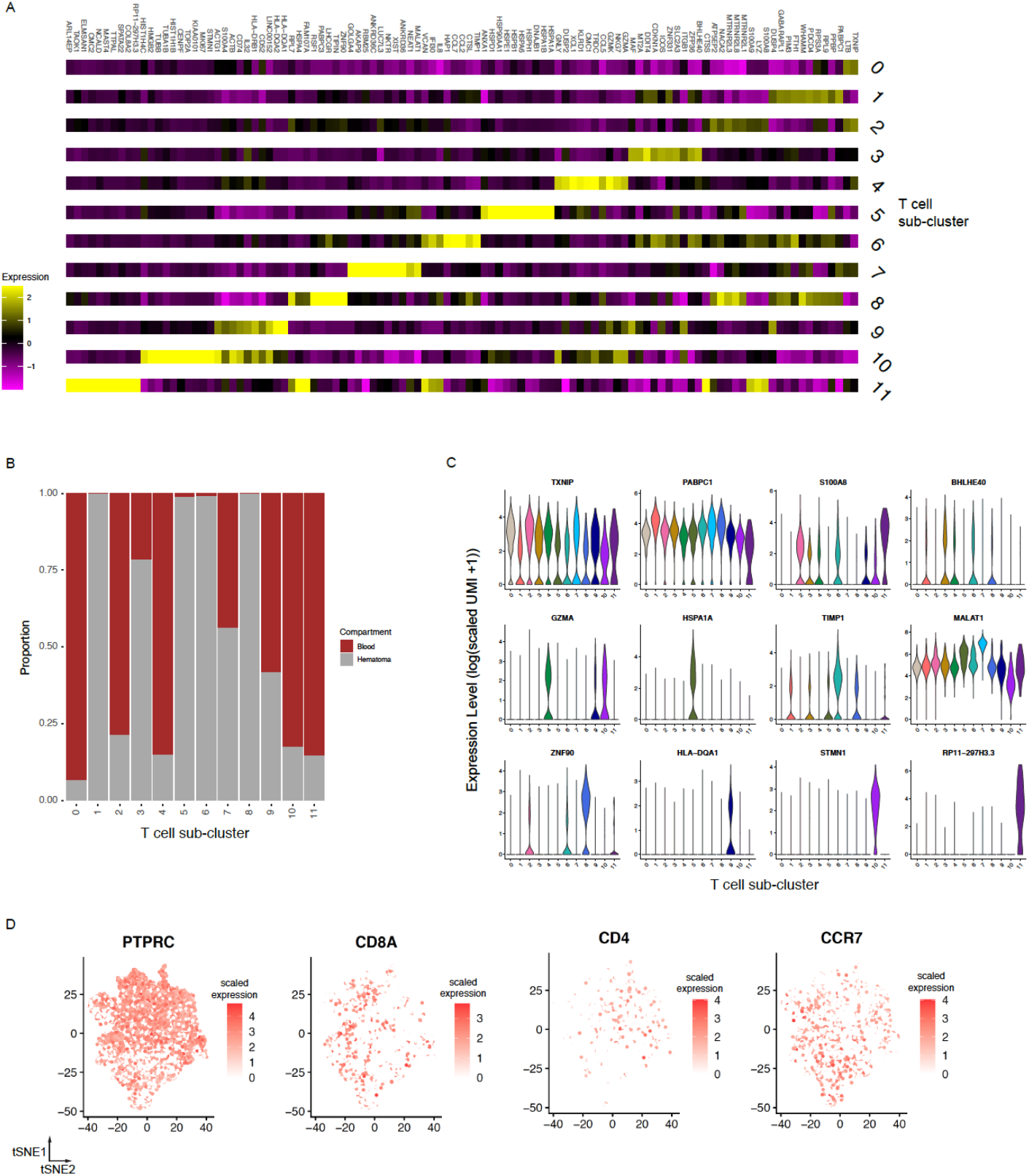
T cell re-clustering analysis. **A**. Heatmap of top ten differentially expressed genes for each T cell sub-cluster. **B**. Stacked bar chart of all T cell sub-clusters by blood or hematoma. **C**. Violin plots of selected marker genes for each T cell sub-cluster. **D**. tSNEs of re-clustered T cell data colored by PTPRC, CD8A, CD4, and CCR7 expression.

**Supplemental Figure 10.**
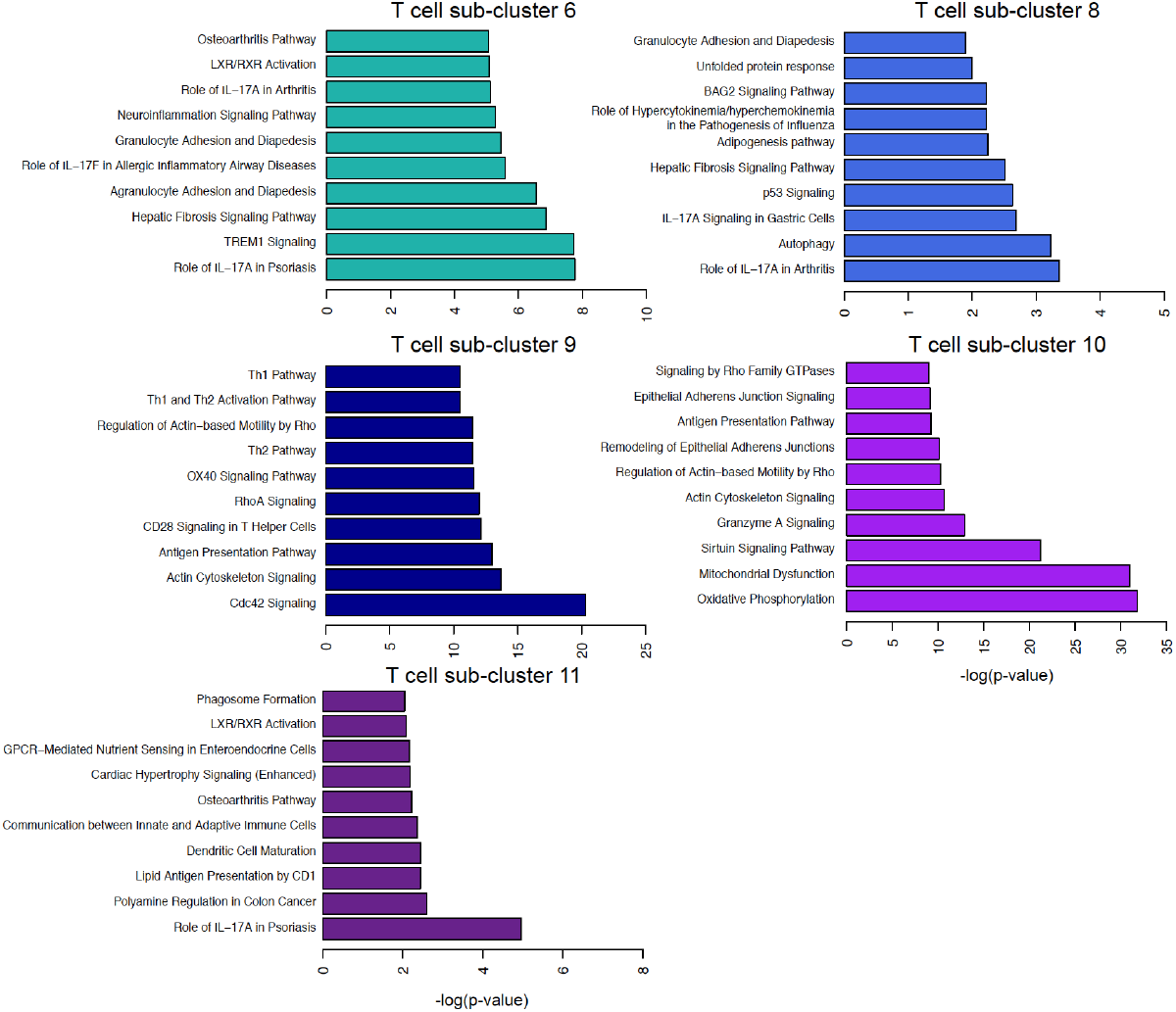
Waterfall plots of top 10 enriched pathways by IPA for remaining T cell sub-clusters. Results, including genes in each pathway, are also shown in Table S9.

**Supplemental Figure 11.**
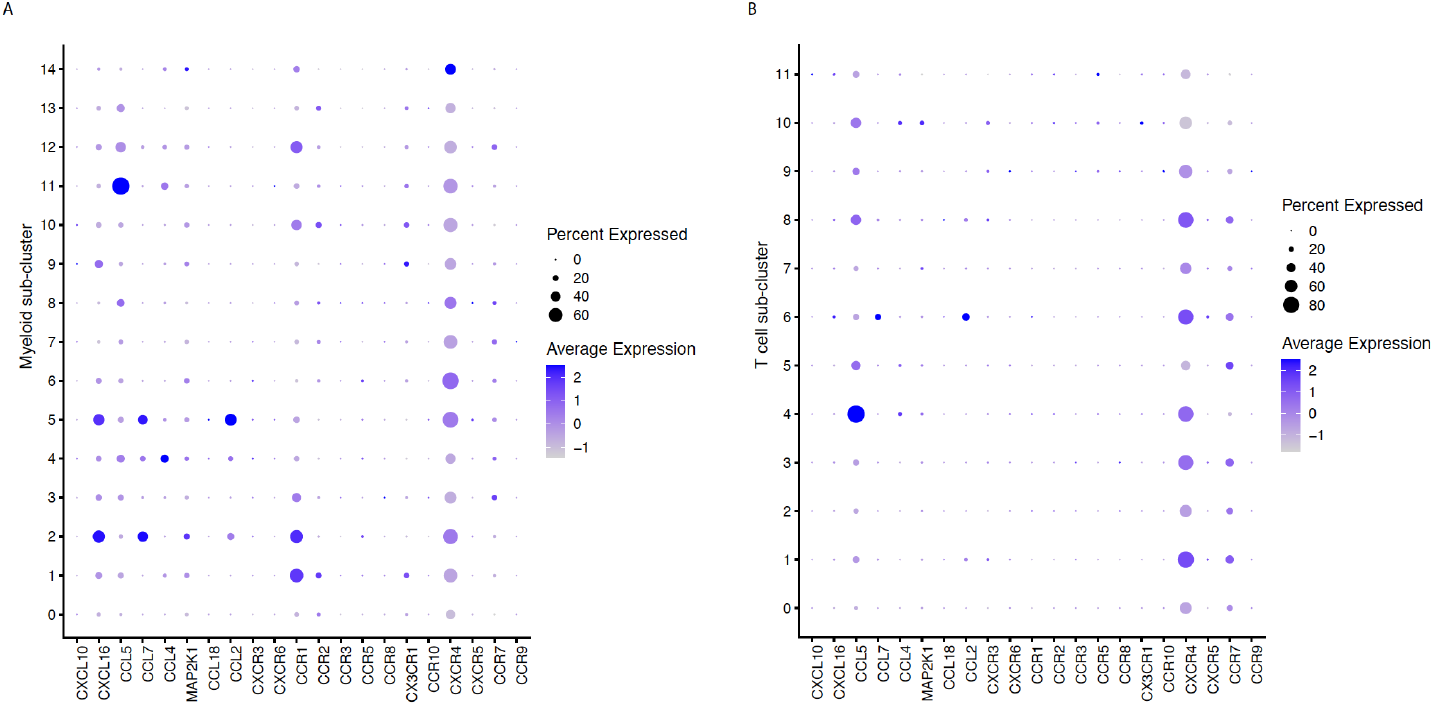
Dot plots of canonical ligands and receptors for **A.** myeloid sub-clusters, and **B.** T cell subclusters.

**Supplemental Figure 12.**
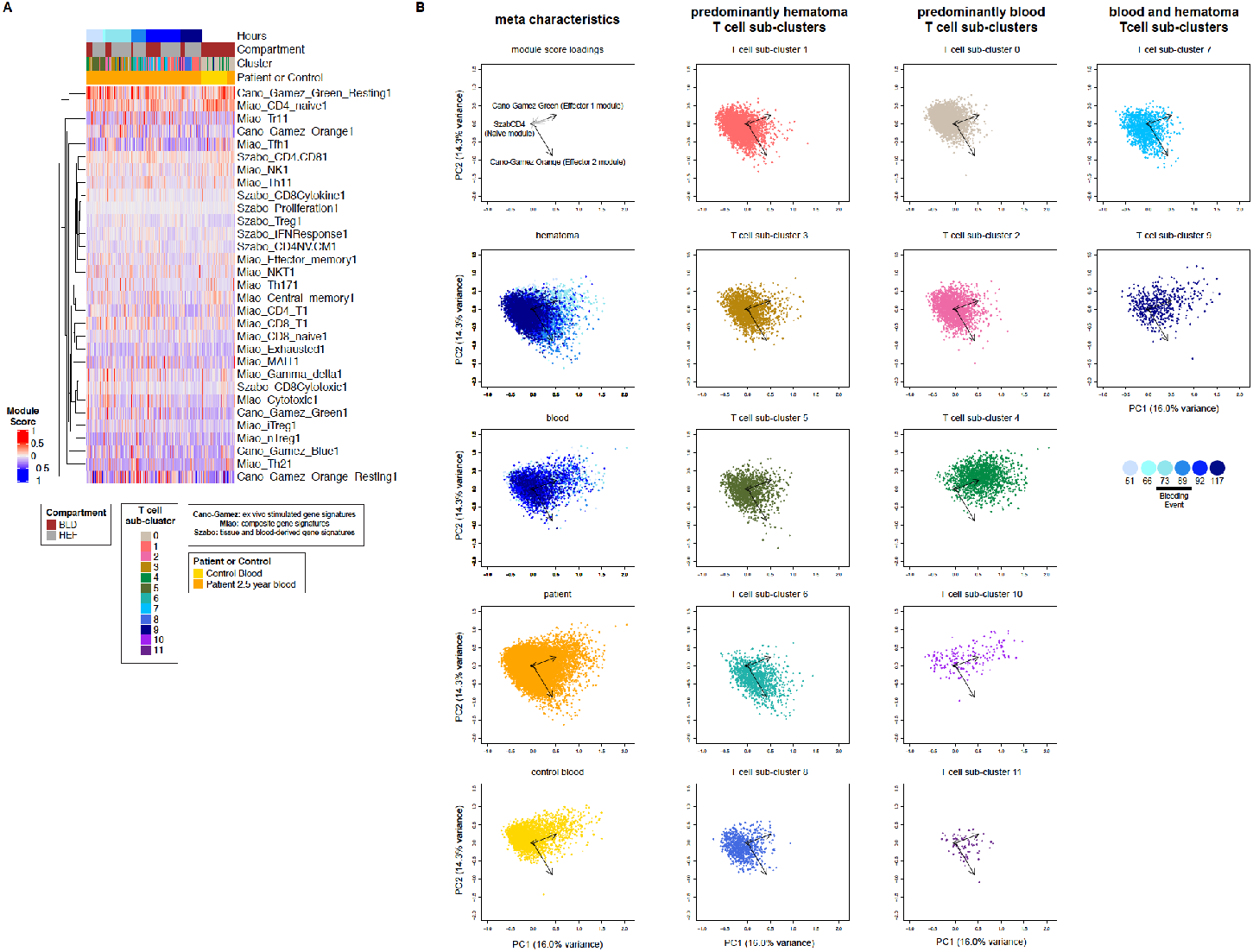
**A**. Gene expression module scores were calculated for selected gene modules from the literature for each cell in our myeloid data and clustered hierarchically. The heatmap is annotated by compartment (blood or hematoma), time, source (control or patient), and T cell sub-cluster. **B**. Principal component analysis was performed on all module scores. Data are shown across the first two principal components and all module variable loadings are projected onto the top left plot and each subsequent plot shows only the top three module loading (black arrows). Data from each myeloid sub-cluster is visualized separately (sub-clusters 0-11), or as a function of time in either blood or hematoma, and as patient derived or control blood derived. Plots are grouped by metadata characteristics, predominantly blood, predominantly hematoma, or both. Gene signatures used for module scoring are described in **Table S10**.

### Supplemental Tables

**Table S1.**
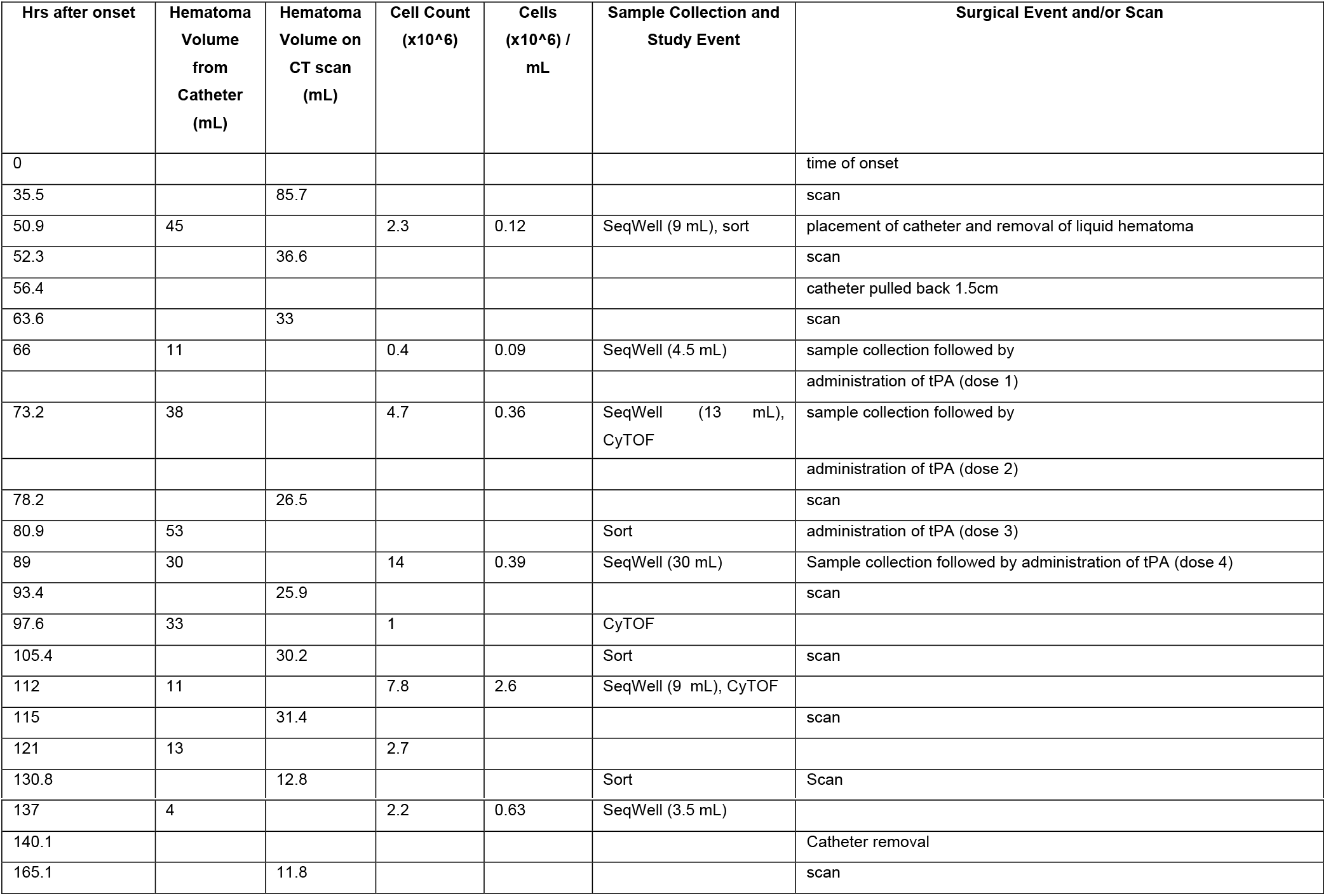
Patient time course.

**Table S2.** CyTOF antibodies.

| Isotope | Epitope | Fluidigm Catalog # |
| --- | --- | --- |
| 89Y | CD45 | 3089003B |
| 142Nd | CD19 | 3142001B |
| 145Nd | CD4 | 3145001B |
| 146Nd | CD8 alpha | 3146001B |
| 147Sm | CD11c | 3147008B |
| 149Sm | CD66a | 3149008B |
| 151Eu | CD14 | 3151009B |
| 167Er | CD11b | 3167011B |
| 170Er | CD3 | 3170001B |
| 174Yb | HLA-DR | 3174001B |
| 176Yb | CD56 | 3176008B |
| 191Ir | DNA (Singlets) | 201192A |
| 193Ir | DNA (Singlets) | 201192A |
| 194Pt | Live/Dead | 201194 |

**Table S3.** scRNA-seq cell counts shown in **Figure 1**.

| Condition (Compartment, Time) | B cells | CD1C+ DCs | CD8 T cells | granulocytes | macrophage | monocyte/macrophage | neurons | NK cells | T cells | TOTAL |
| --- | --- | --- | --- | --- | --- | --- | --- | --- | --- | --- |
| Blood, 51 hrs | 33 | 72 | 251 | 27 | 20 | 834 | 0 | 62 | 483 | 1782 |
| Blood, 66 hrs | 18 | 10 | 0 | 15 | 13 | 181 | 0 | 11 | 57 | 305 |
| Blood, 73 hrs | 38 | 4 | 100 | 7 | 36 | 323 | 0 | 278 | 664 | 1450 |
| Blood, 89 hrs | 57 | 22 | 110 | 20 | 198 | 872 | 0 | 209 | 1543 | 3031 |
| Blood, 112 hrs | 49 | 26 | 147 | 13 | 40 | 158 | 0 | 358 | 1590 | 2381 |
| Blood, 137 hrs | 9 | 13 | 10 | 13 | 61 | 165 | 0 | 59 | 484 | 814 |
| Blood, patient followup | 133 | 88 | 70 | 20 | 1 | 327 | 0 | 197 | 794 | 1630 |
| Blood, control 00 | 67 | 13 | 77 | 2 | 0 | 47 | 0 | 425 | 1184 | 1815 |
| Blood, control 01 | 35 | 74 | 29 | 27 | 9 | 771 | 0 | 150 | 518 | 1613 |
| Blood, control 02 | 10 | 60 | 4 | 15 | 5 | 1531 | 0 | 21 | 151 | 1797 |
| Blood, control 03 | 56 | 56 | 192 | 9 | 0 | 313 | 0 | 238 | 756 | 1620 |
| Hematoma, 51 hrs | 13 | 2 | 9 | 1 | 425 | 0 | 17 | 18 | 1126 | 1611 |
| Hematoma, 66 hrs | 2 | 0 | 0 | 0 | 12 | 0 | 351 | 6 | 248 | 619 |
| Hematoma, 73 hrs | 91 | 55 | 200 | 20 | 816 | 39 | 1 | 278 | 2145 | 3645 |
| Hematoma, 89 hrs | 102 | 9 | 20 | 9 | 121 | 3 | 0 | 171 | 1422 | 1857 |
| Hematoma, 112 hrs | 83 | 17 | 12 | 6 | 235 | 13 | 8 | 601 | 2336 | 3311 |
| Hematoma, 137 hrs | 32 | 4 | 1 | 2 | 196 | 2 | 7 | 307 | 1890 | 2441 |

**Table S4.**
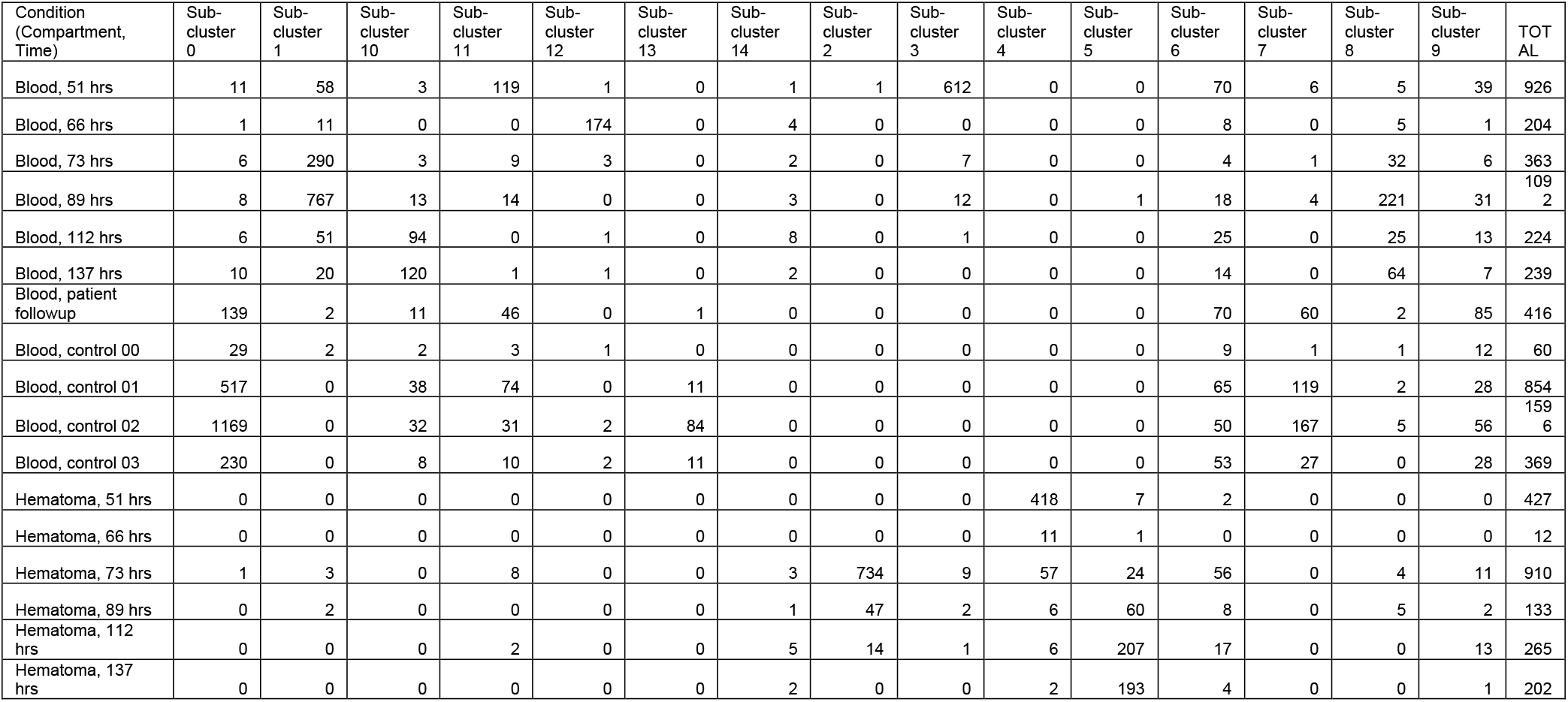
Myeloid cell re-clustering analysis cluster membership cell counts.

**Table S5.**
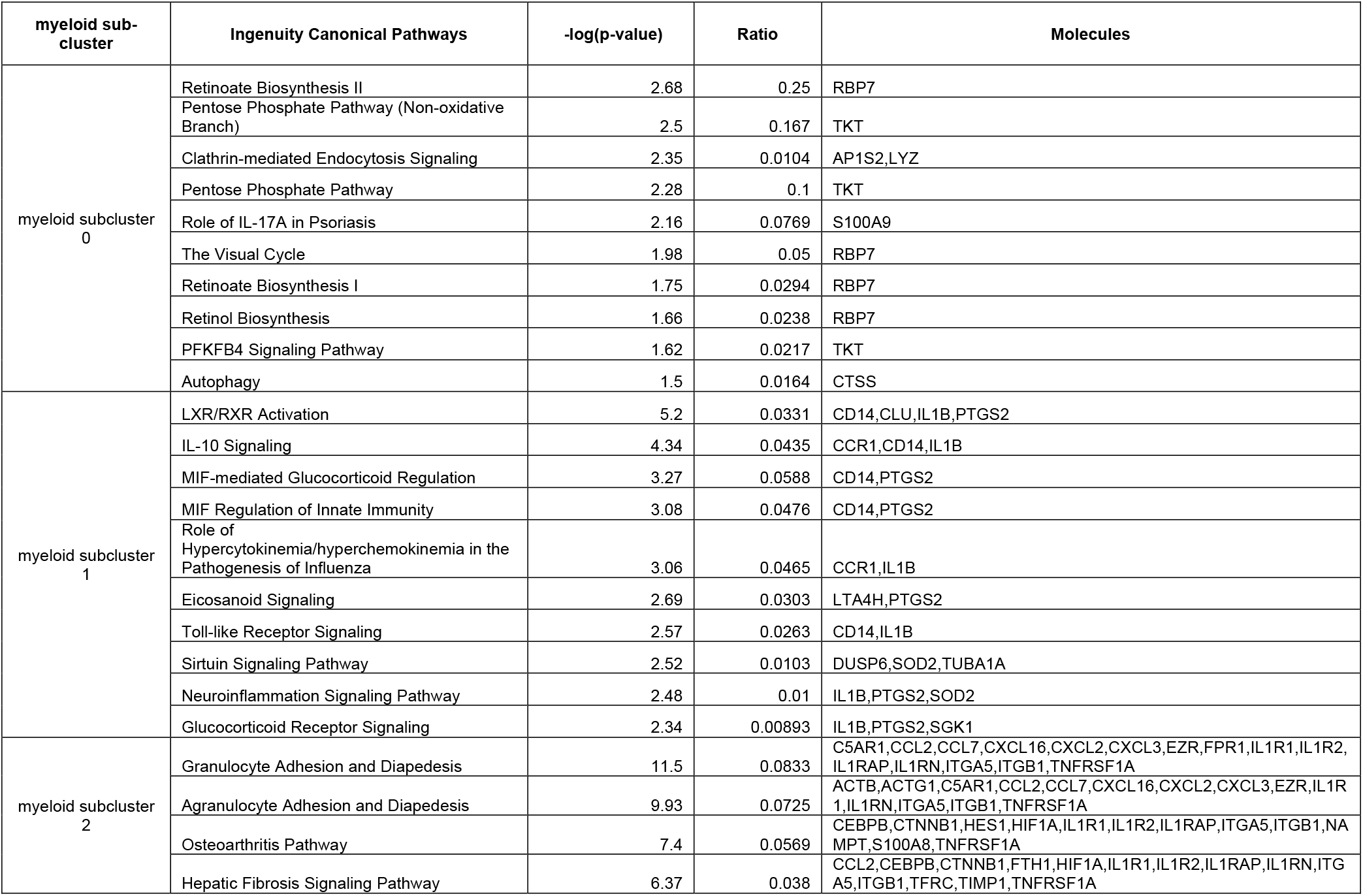
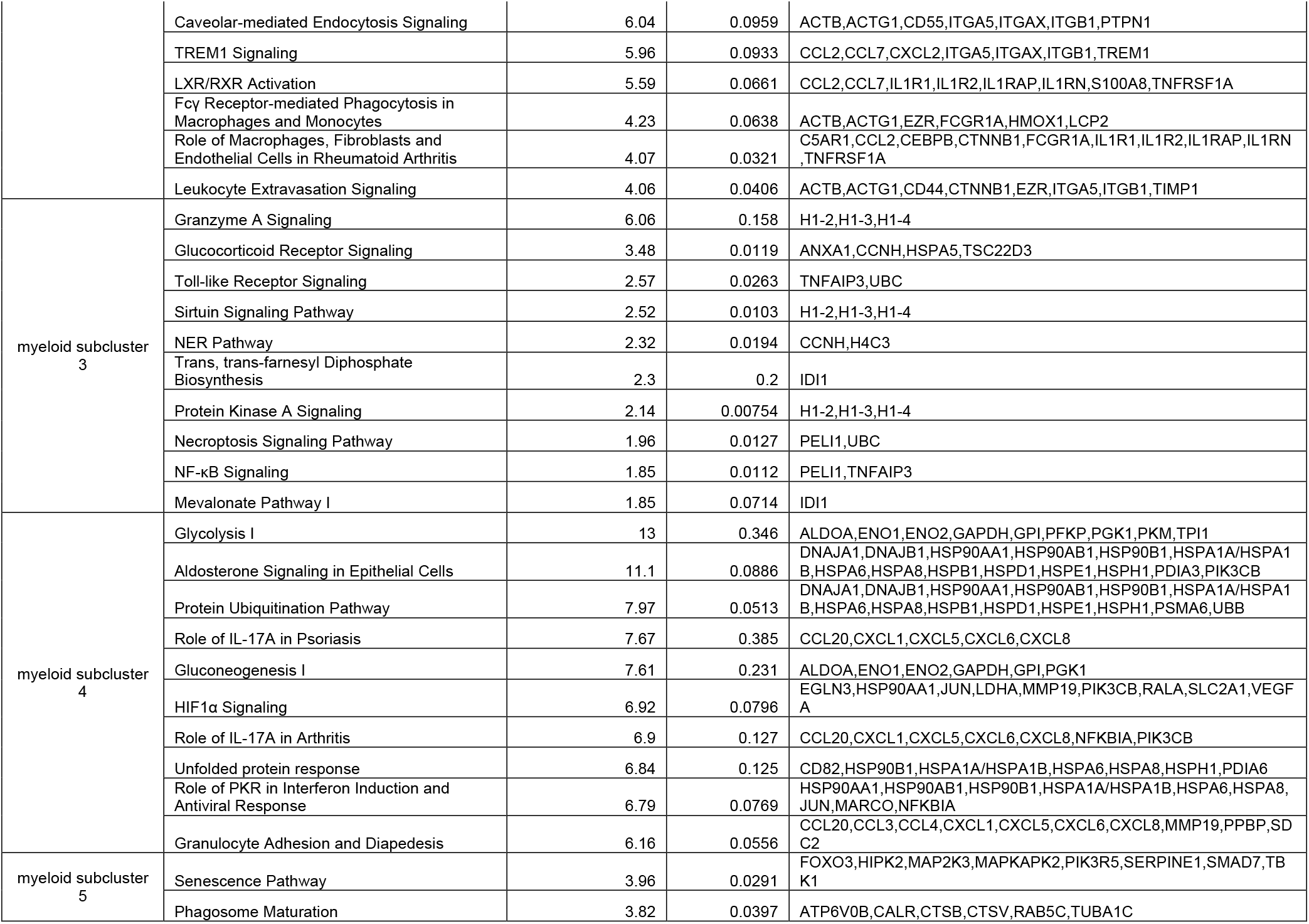
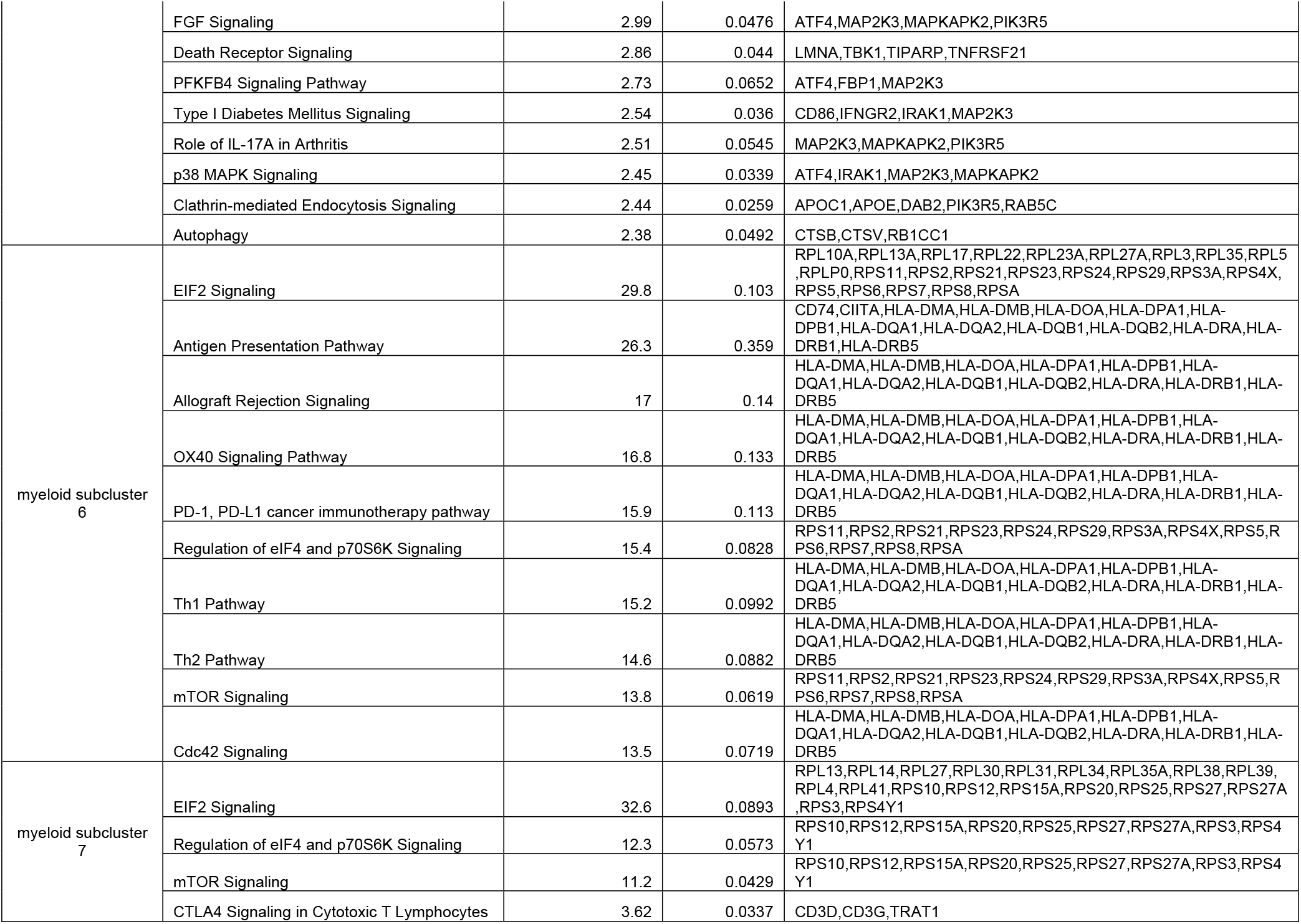
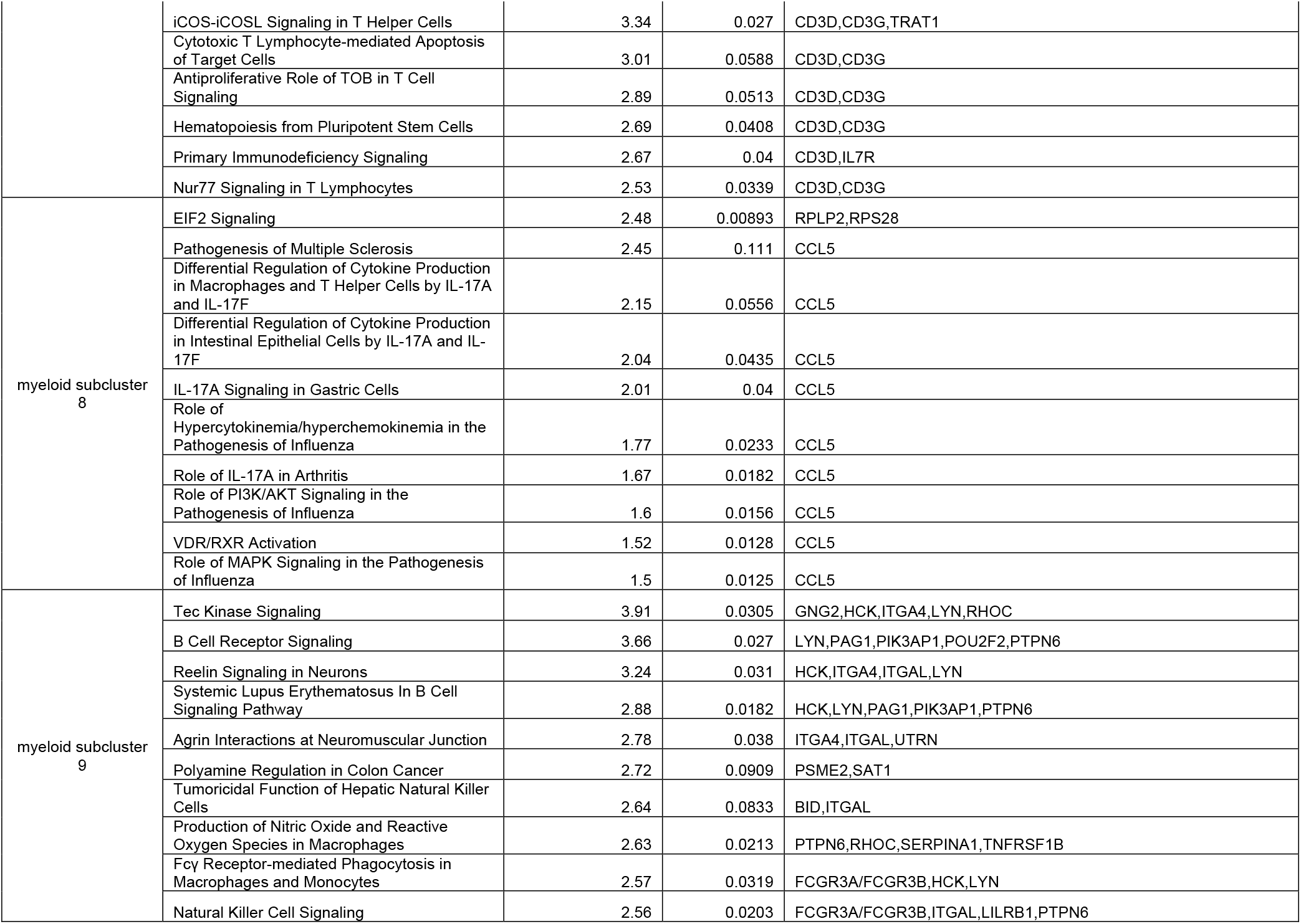
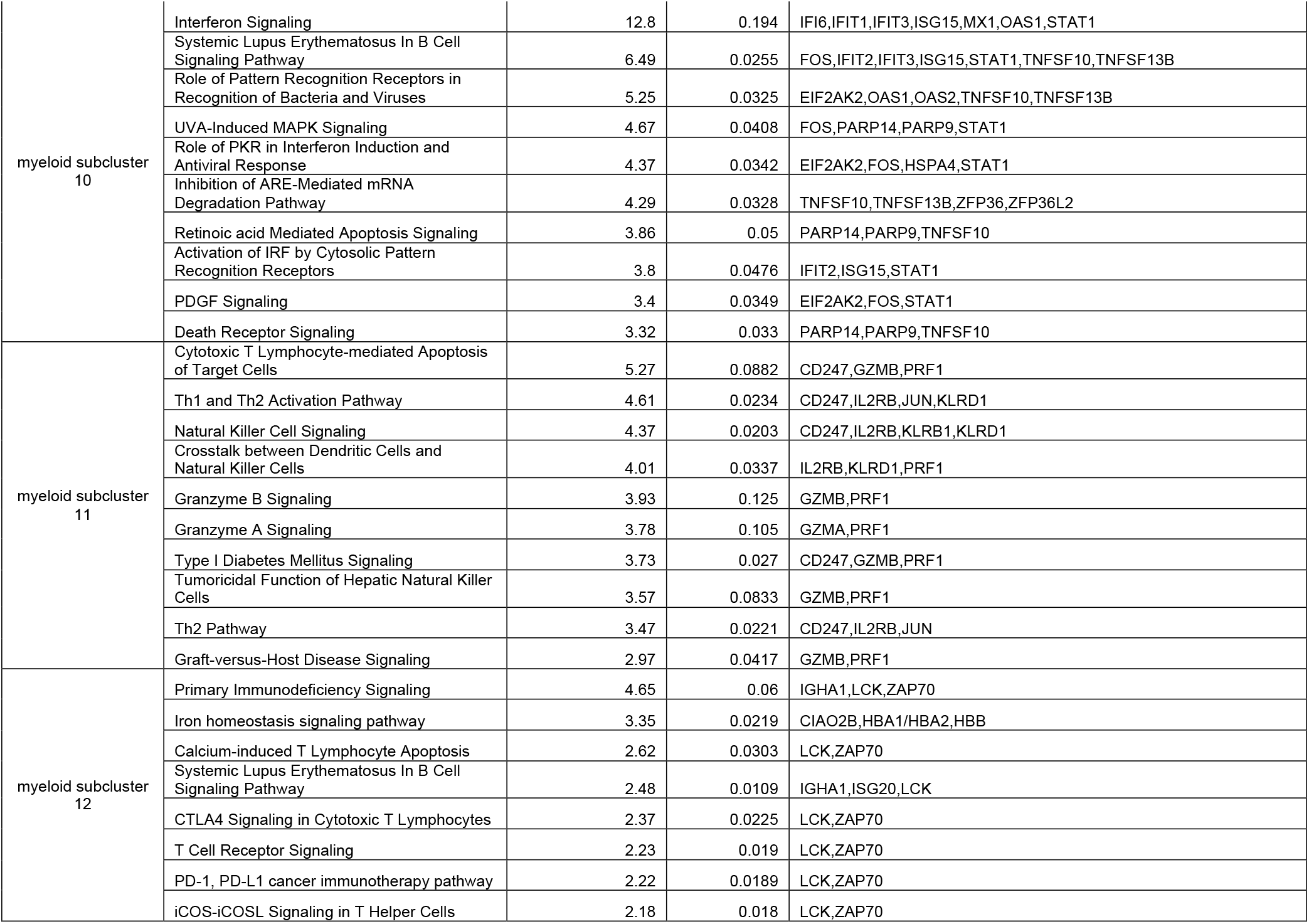
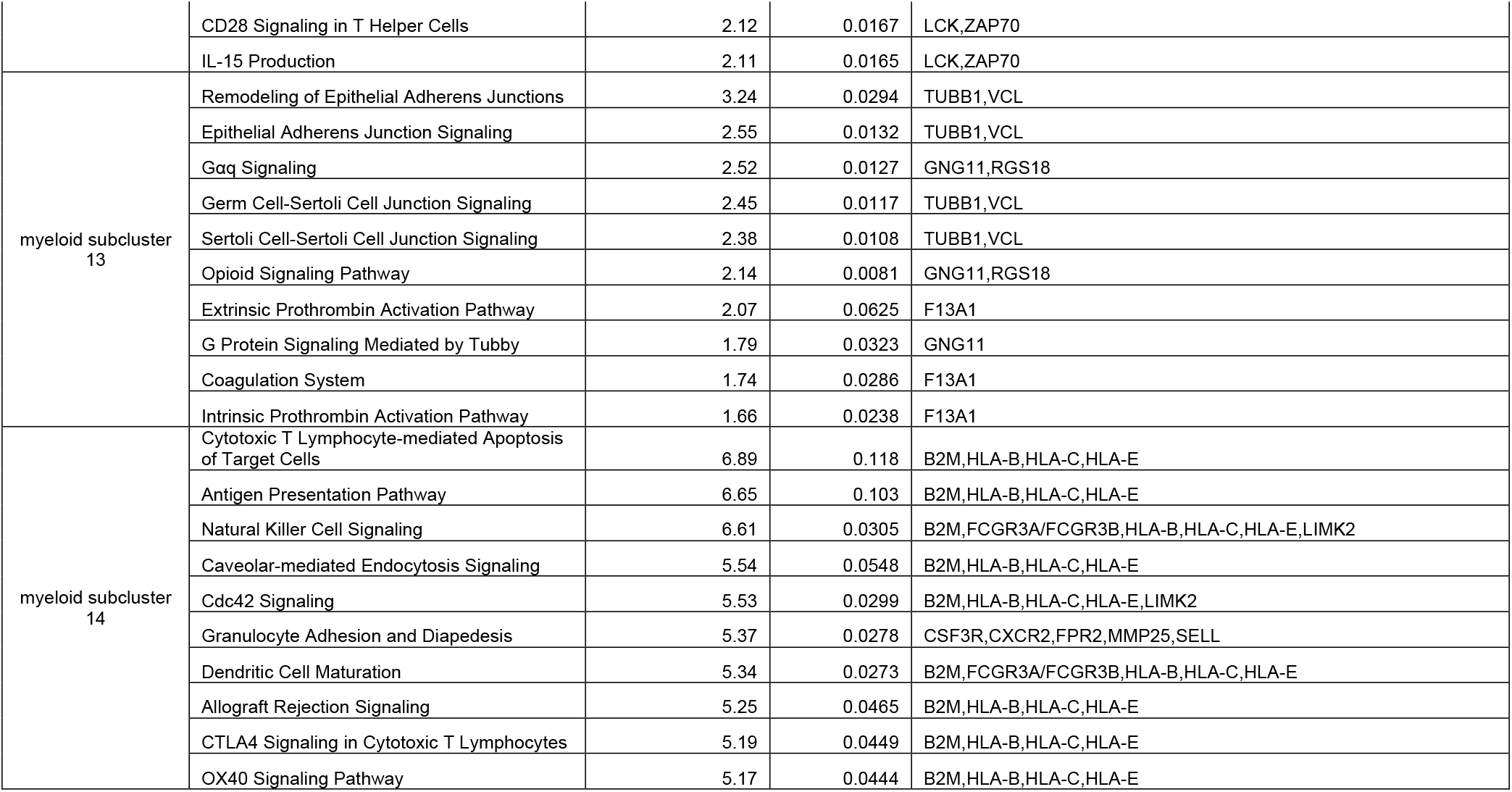
Monocyte sub-cluster IPA results. Top 10 pathways and associated genes are shown.

**Table S6.** Human monocyte gene sets used for module scoring.

| **Gene Set Name** | **Description** | **Reference** |
| --- | --- | --- |
| CD141_CLEC9A_Villani | Generated from unstimulated human PBMCs, identified by scRNA-seq. | Villani et al.<sup>3</sup> |
| CD1C_A_Villani | Generated from unstimulated human PBMCs, identified by scRNA-seq. | Villani et al.<sup>3</sup> |
| CD1C_B_Villani | Generated from unstimulated human PBMCs, identified by scRNA-seq. | Villani et al.<sup>3</sup> |
| CD1Cminus_CD141minus_Villani | Generated from unstimulated human PBMCs, identified by scRNA-seq. | Villani et al.<sup>3</sup> |
| New_pop_Villani | Generated from unstimulated human PBMCs, identified by scRNA-seq. | Villani et al.<sup>3</sup> |
| pDC_Villani | Generated from unstimulated human PBMCs, identified by scRNA-seq. | Villani et al.<sup>3</sup> |
| XUE_Module1 to XUE_Module49 | Generated by comparative/ integrative analysis on several microarray datasets. | Xue et al.<sup>3</sup> |
| Monocyte_human | ImmGenn database | Shay et al.<sup>2</sup> |
| MyeloidDC_human | ImmGenn database | Shay et al.<sup>2</sup> |
| PlasmacytoidDC_human | ImmGenn database | Shay et al.<sup>2</sup> |

**Table S7.** Wilcox rank sum pairwise test results for CD1CB (CD14^+^ monocyte) signature module scores, data presented in Figure 2. Table displays adjusted p values (p_adj_) for pairwise comparisons for each indicated myeloid subcluster. P values are colored as follows: blue (0.01 ≤ p_adj_ < 0.05, *), purple (0.001 ≤ p_adj_ < 0.01, **), white (0 ≤ p_adj_ < 0.001, ***), gray (not significant, padj < 0.05).

| Sub-cluster | 0 | 1 | 2 | 3 | 4 | 5 | 6 | 7 | 8 | 9 | 10 | 11 | 12 | 13 |
| --- | --- | --- | --- | --- | --- | --- | --- | --- | --- | --- | --- | --- | --- | --- |
| 1 | 1.68E-47 | - | - | - | - | - | - | - | - | - | - | - | - | - |
| 2 | 6.19E-71 | 3.28E-10 | - | - | - | - | - | - | - | - | - | - | - | - |
| 3 | 1.15E-16 | 1.72E-57 | 1.66E-69 | - | - | - | - | - | - | - | - | - | - | - |
| 4 | 3.08E-249 | 3.56E-225 | 1.74E-193 | 1.74E-152 | - | - | - | - | - | - | - | - | - | - |
| 5 | 2.06E-224 | 2.56E-212 | 3.16E-183 | 5.98E-125 | 2.91E-08 | - | - | - | - | - | - | - | - | - |
| 6 | 2.95E-187 | 7.73E-189 | 1.50E-166 | 6.25E-98 | 2.16E-06 | 6.31E-01 | - | - | - | - | - | - | - | - |
| 7 | 1.03E-65 | 8.67E-103 | 2.51E-100 | 1.06E-14 | 4.04E-106 | 7.82E-78 | 2.00E-54 | - | - | - | - | - | - | - |
| 8 | 6.51E-92 | 6.62E-120 | 4.97E-112 | 2.32E-30 | 3.11E-92 | 1.19E-60 | 3.87E-38 | 4.82E-06 | - | - | - | - | - | - |
| 9 | 4.48E-184 | 3.41E-169 | 1.08E-151 | 9.39E-134 | 3.03E-38 | 1.19E-52 | 1.00E-37 | 1.35E-104 | 1.01E-98 | - | - | - | - | - |
| 10 | 1.29E-02 | 6.82E-24 | 1.02E-35 | 5.30E-04 | 1.66E-117 | 1.91E-103 | 4.21E-86 | 6.68E-26 | 3.08E-41 | 3.85E-104 | - | - | - | - |
| 11 | 1.80E-39 | 2.42E-71 | 4.43E-75 | 5.83E-09 | 1.75E-72 | 6.97E-55 | 3.53E-41 | 3.48E-01 | 3.85E-06 | 1.62E-78 | 1.29E-16 | - | - | - |
| 12 | 2.78E-02 | 1.59E-17 | 4.05E-26 | 3.21E-03 | 1.69E-83 | 1.16E-74 | 9.99E-63 | 6.67E-21 | 1.16E-32 | 1.11E-76 | 9.52E-01 | 1.63E-13 | - | - |
| 13 | 7.50E-04 | 4.59E-15 | 1.94E-20 | 4.75E-01 | 2.89E-52 | 1.81E-45 | 6.63E-37 | 5.65E-09 | 9.59E-17 | 1.69E-51 | 1.14E-01 | 6.67E-06 | 1.23E-01 | - |
| 14 | 4.48E-08 | 1.68E-12 | 2.35E-13 | 1.57E-03 | 1.35E-14 | 1.28E-10 | 1.04E-07 | 4.92E-01 | 3.47E-01 | 1.00E-17 | 4.53E-06 | 3.72E-01 | 3.39E-06 | 3.69E-04 |

**Table S8.** T cell sub-clustering analysis cluster membership cell counts. Sub-clusters are indicated by NT0-NT11.

| Condition (Compartment, Time) | NT 0 | NT 1 | NT 2 | NT 3 | NT 4 | NT 5 | NT 6 | NT 7 | NT 8 | NT 9 | NT 10 | NT 11 | Total |
| --- | --- | --- | --- | --- | --- | --- | --- | --- | --- | --- | --- | --- | --- |
| Blood, 51 hrs | 34 | 1 | 10 | 323 | 268 | 17 | 0 | 1 | 0 | 52 | 15 | 0 | 721 |
| Blood, 66 hrs | 1 | 0 | 17 | 0 | 3 | 0 | 3 | 4 | 0 | 0 | 0 | 0 | 28 |
| Blood, 73 hrs | 27 | 0 | 485 | 21 | 118 | 0 | 0 | 14 | 0 | 22 | 8 | 0 | 695 |
| Blood, 89 hrs | 3 | 0 | 49 | 1 | 101 | 0 | 1 | 8 | 0 | 10 | 11 | 0 | 184 |
| Blood, 112 hrs | 326 | 0 | 579 | 8 | 237 | 0 | 3 | 456 | 0 | 55 | 12 | 5 | 1681 |
| Blood, 137 hrs | 4 | 0 | 365 | 0 | 14 | 0 | 6 | 22 | 1 | 2 | 0 | 53 | 467 |
| Blood, patient followup | 645 | 0 | 6 | 6 | 95 | 0 | 0 | 0 | 0 | 58 | 48 | 0 | 858 |
| Blood, control 00 | 1069 | 1 | 1 | 16 | 109 | 0 | 0 | 2 | 0 | 45 | 17 | 0 | 1260 |
| Blood, control 01 | 424 | 0 | 4 | 1 | 61 | 0 | 0 | 0 | 0 | 18 | 15 | 0 | 523 |
| Blood, control 02 | 133 | 0 | 2 | 3 | 5 | 0 | 0 | 0 | 0 | 3 | 3 | 0 | 149 |
| Blood, control 03 | 555 | 1 | 11 | 6 | 325 | 0 | 0 | 4 | 0 | 29 | 8 | 0 | 939 |
| Hematoma, 51 hrs | 4 | 4 | 2 | 18 | 8 | 1087 | 1 | 11 | 0 | 0 | 0 | 0 | 1135 |
| Hematoma, 66 hrs | 3 | 7 | 3 | 1 | 0 | 217 | 4 | 11 | 2 | 0 | 0 | 0 | 248 |
| Hematoma, 73 hrs | 194 | 37 | 345 | 1266 | 190 | 19 | 81 | 56 | 0 | 133 | 11 | 0 | 2332 |
| Hematoma, 89 hrs | 2 | 116 | 54 | 66 | 23 | 4 | 1014 | 122 | 4 | 15 | 4 | 0 | 1424 |
| Hematoma, 112 hrs | 25 | 1774 | 6 | 45 | 10 | 1 | 43 | 355 | 18 | 54 | 12 | 5 | 2348 |
| Hematoma, 137 hrs | 2 | 1064 | 2 | 0 | 0 | 5 | 12 | 98 | 692 | 9 | 2 | 5 | 1891 |

**Table S9.**
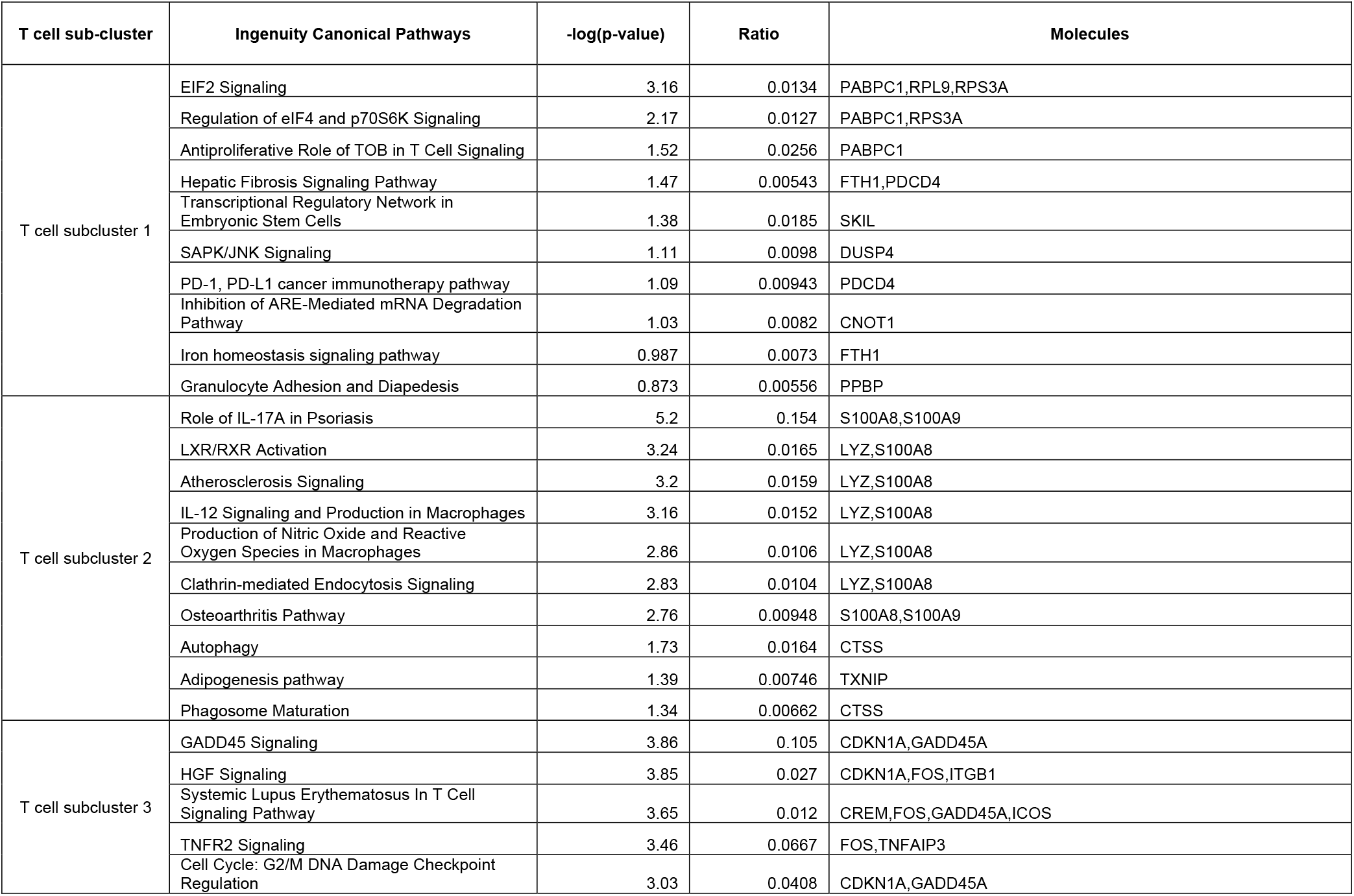
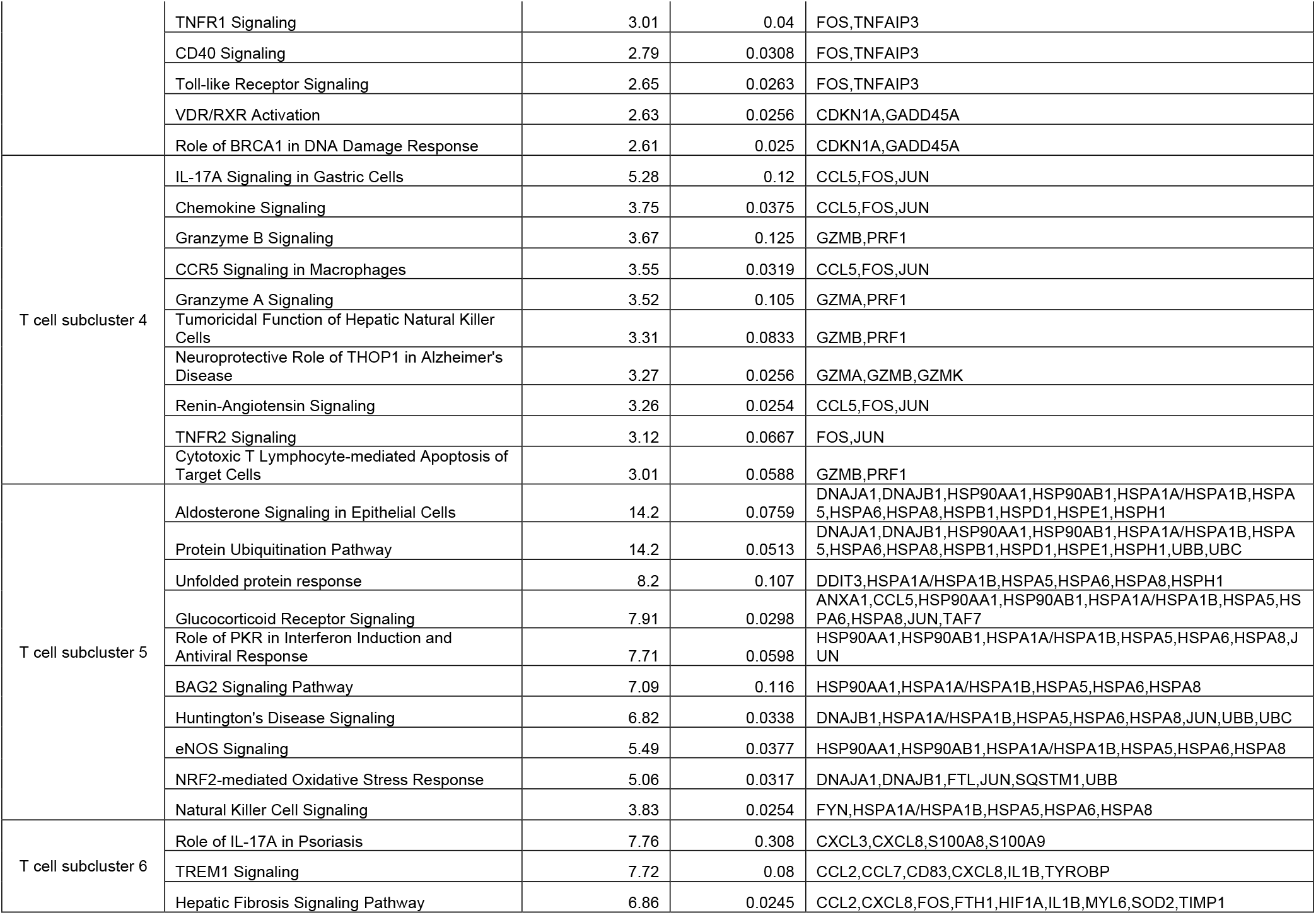
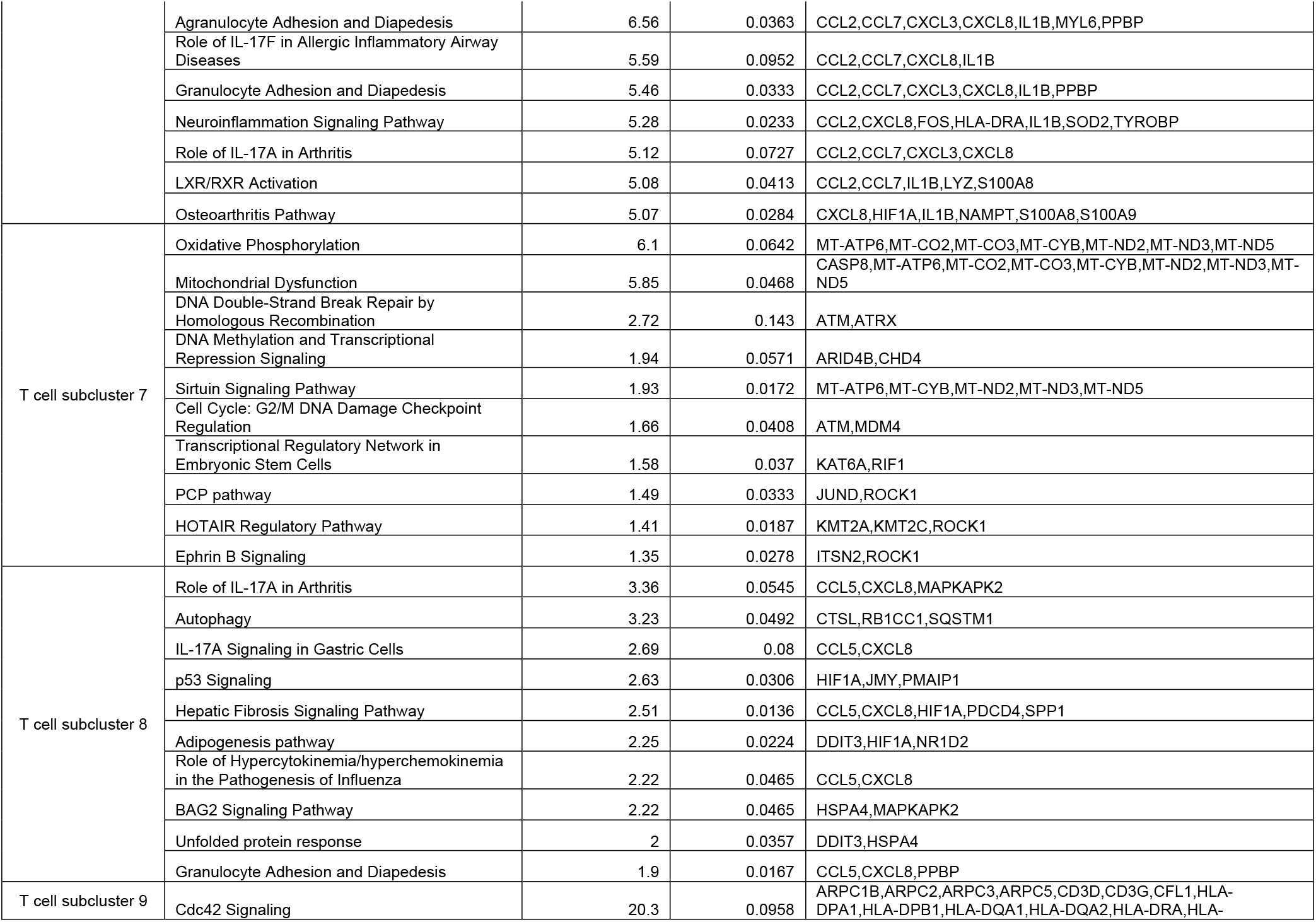
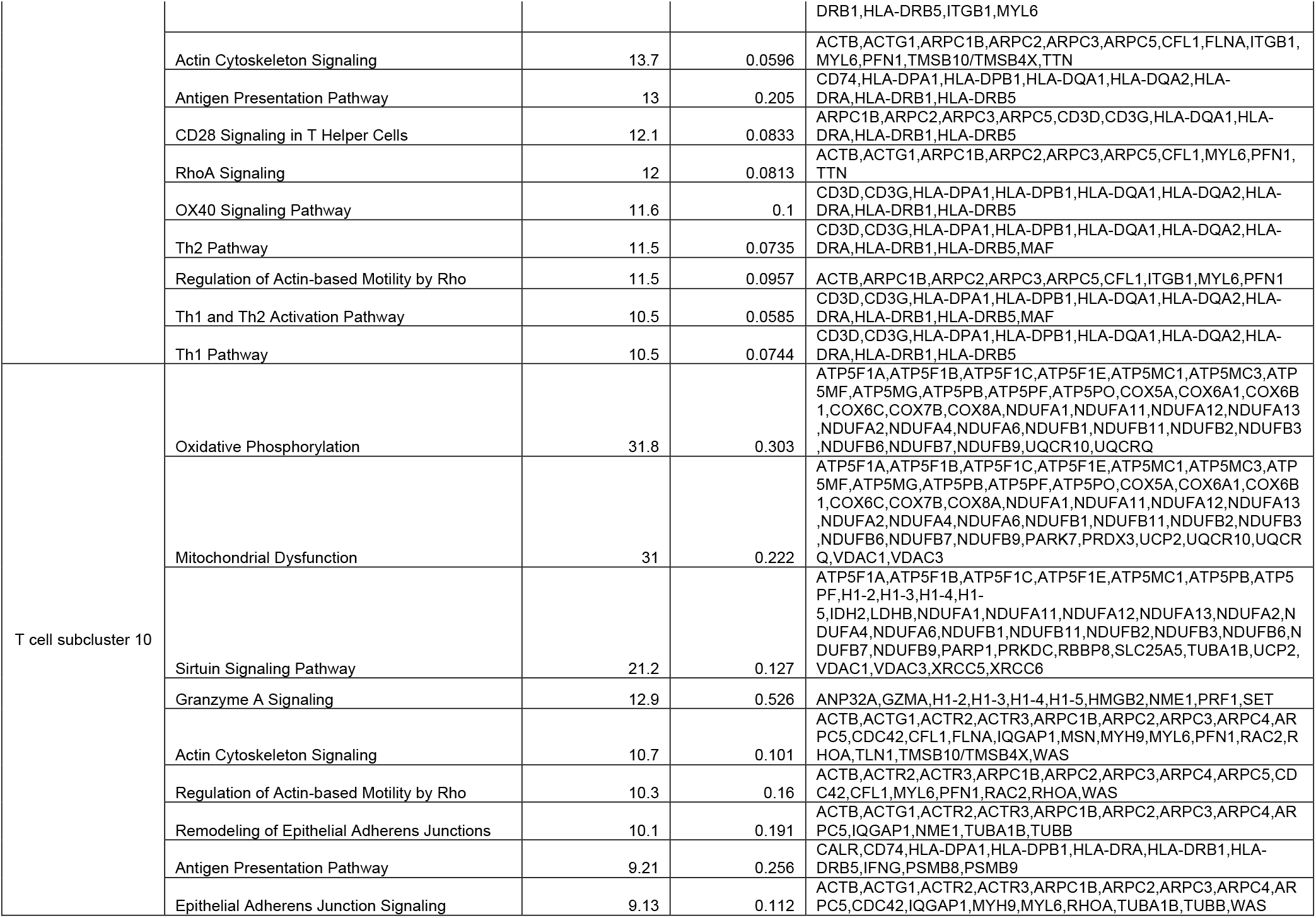
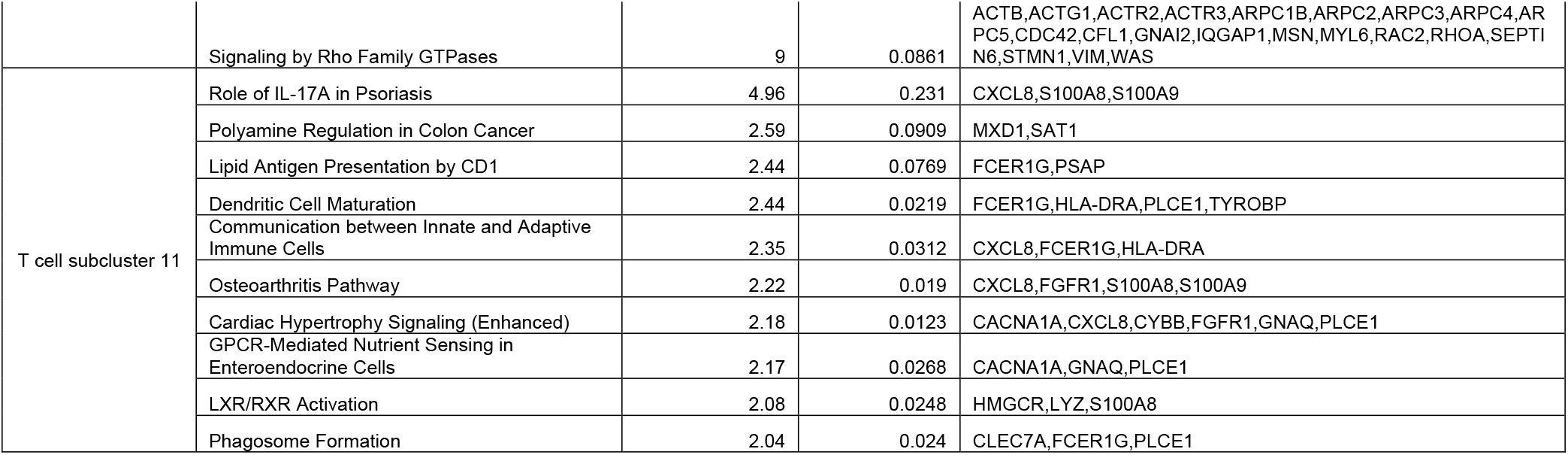
T cell sub-cluster IPA results. Top 10 pathways and associated genes are shown.

**Table S10.** Human monocyte gene sets used for module scoring.

| **Gene Set Name** | **Description** | **Reference** |
| --- | --- | --- |
| Szabo gene sets | Generated from resting and stimulated human T cells isolated from blood and tissues (lungs, lymph nodes, bone marrow). | Szabo et al.<sup>6</sup> |
| Miao gene sets | Generated from summary of marker genes across the literature (Table S1). | Miao et al.<sup>7</sup> |
| Cano-Gamez gene sets | Generated from unstimulated and stimulated human T cells, identified by scRNA-seq. | Cano-Gamez et al.<sup>8</sup> |

## Notes

### Competing Interest Statement

The authors have declared no competing interest.

